# Feature-based Molecular Networking in the GNPS Analysis Environment

**DOI:** 10.1101/812404

**Authors:** Louis Felix Nothias, Daniel Petras, Robin Schmid, Kai Dührkop, Johannes Rainer, Abinesh Sarvepalli, Ivan Protsyuk, Madeleine Ernst, Hiroshi Tsugawa, Markus Fleischauer, Fabian Aicheler, Alexander Aksenov, Oliver Alka, Pierre-Marie Allard, Aiko Barsch, Xavier Cachet, Mauricio Caraballo, Ricardo R. Da Silva, Tam Dang, Neha Garg, Julia M. Gauglitz, Alexey Gurevich, Giorgis Isaac, Alan K. Jarmusch, Zdeněk Kameník, Kyo Bin Kang, Nikolas Kessler, Irina Koester, Ansgar Korf, Audrey Le Gouellec, Marcus Ludwig, Martin H. Christian, Laura-Isobel McCall, Jonathan McSayles, Sven W. Meyer, Hosein Mohimani, Mustafa Morsy, Oriane Moyne, Steffen Neumann, Heiko Neuweger, Ngoc Hung Nguyen, Melissa Nothias-Esposito, Julien Paolini, Vanessa V. Phelan, Tomáš Pluskal, Robert A. Quinn, Simon Rogers, Bindesh Shrestha, Anupriya Tripathi, Justin J.J. van der Hooft, Fernando Vargas, Kelly C. Weldon, Michael Witting, Heejung Yang, Zheng Zhang, Florian Zubeil, Oliver Kohlbacher, Sebastian Böcker, Theodore Alexandrov, Nuno Bandeira, Mingxun Wang, Pieter C. Dorrestein

## Abstract

Molecular networking has become a key method used to visualize and annotate the chemical space in non-targeted mass spectrometry-based experiments. However, distinguishing isomeric compounds and quantitative interpretation are currently limited. Therefore, we created Feature-based Molecular Networking (FBMN) as a new analysis method in the Global Natural Products Social Molecular Networking (GNPS) infrastructure. FBMN leverages feature detection and alignment tools to enhance quantitative analyses and isomer distinction, including from ion-mobility spectrometry experiments, in molecular networks.

## Main text

Introduced in 2012^1^, molecular networking has become an essential bioinformatics tool to visualize and annotate non-targeted mass spectrometry data^2,3^. The first application of molecular networking was described by Traxler and Kolter^4^ as holding “*great promise in providing the next quantum leap in understanding the fascinating world of microbial chemical ecology*”. Molecular networking goes beyond spectral matching against reference spectra, by aligning experimental spectra against one another and connecting related molecules by their spectral similarity^5^. In a molecular network, related molecules are referred to as a “molecular family”, differing by simple transformations such as glycosylation, alkylation, and oxidation/reduction. Molecular networking became publicly accessible in 2013 through the initial release of the Global Natural Product Social Molecular Networking (GNPS), a web-enabled mass spectrometry knowledge capture and analysis platform (http://gnps.ucsd.edu)^6^, and has been widely applied in mass spectrometry-based metabolomics to aid in the annotation of molecular families from their fragmentation spectra (MS^2^).

Powered by 3,000+ CPU cores at the Center for Computational Mass Spectrometry at the University of California San Diego and the MassIVE data repository, GNPS has provided researchers from more than 150 countries with the ability to perform molecular networking. To build upon the success of the first molecular networking method (referred to as “classical” molecular networking, classical MN) which is based on the MS-Cluster algorithm^7^, we introduce a complementary tool named Feature-based Molecular Networking (FBMN). FBMN accepts the outputs of well-established mass spectrometry processing software and improves upon classical MN by incorporating MS^1^ information, such as isotope patterns and retention time, but also ion-mobility separation when performed. As a result, molecular networks obtained with FBMN can distinguish isomers that may have remained hidden, facilitates spectral annotation, and incorporates relative quantitative information which enables robust downstream metabolomics statistical analysis. Whereas users of the classical molecular networking would have had to perform molecular networking and MS^1^ analysis separately before performing a cumbersome linking of the outputs, a key advantage of FBMN is that the analysis is fully integrated throughout the analysis pipeline.

To fully utilize the MS^1^ and MS^2^ content collected during a non-targeted liquid chromatography coupled to tandem mass spectrometry data (LC-MS^2^) metabolomics experiment in a streamlined fashion, we have created an online infrastructure to support the outputs of feature detection and alignment tools for FBMN analysis (https://ccms-ucsd.github.io/GNPSDocumentation/featurebasedmolecularnetworking), including the standard output format for small molecules analysis (mzTab-M)^8^ (Fig. 1a). The diversity of supported software, each offering different functionalities/modules, serves experimentalists, bioinformaticians, and software developers. FBMN is already the second most utilized analysis tool within the GNPS environment (Fig. 1b) with more than 600 jobs performed in August 2019 and has already been used in more than 80 publications using FBMN during its development since Nov 2017. The molecular networks generated with FBMN enable the efficient visualization and annotation of isomers in LC-MS^2^ datasets, as demonstrated below with LC-MS^2^ data from a drug discovery project from *Euphorbia* plant extract^9^ (Fig. 2a-b), and the detection of human microbiome-derived lipids, belonging to the commendamide family^10^, detected in fecal samples from the American Gut Project^11^ (a crowd-sourced citizen science microbiome project) (Fig. 2c-d). In both cases, FBMN resolved positional isomers in the molecular networks that have similar MS^2^ spectra but distinct retention times, that would not have been resolved with classical MN. In the study of metabolites produced by *Euphorbia* plants, the annotation of isomers in the FBMN facilitated the subsequent isolation of antiviral compounds^12^. With samples from the American Gut Project, FBMN enabled the annotation of commendamide isomers, including a putative novel derivative^11^.

**Fig. 1:**
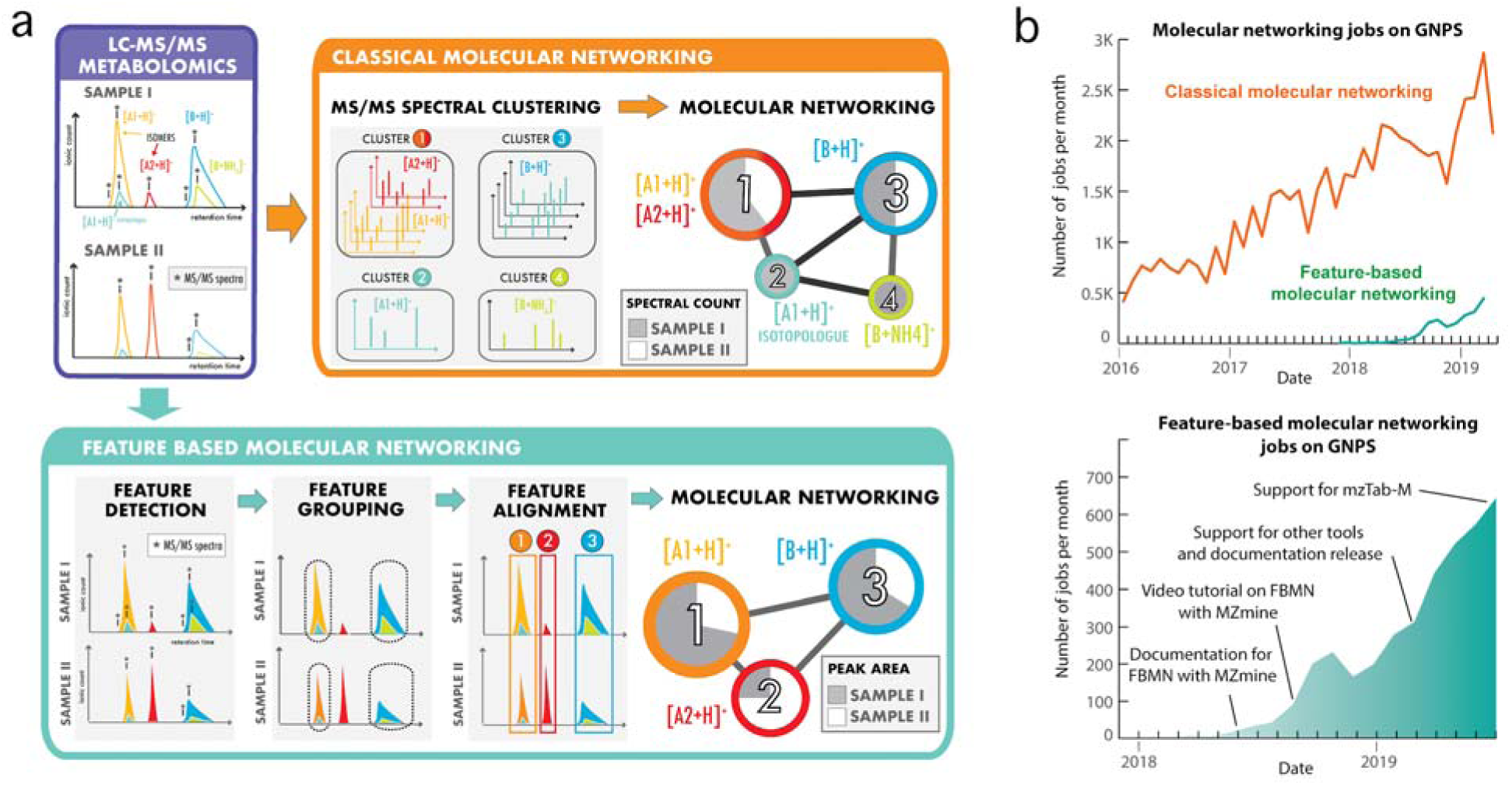
Methods for the generation of molecular networks from non-targeted mass spectrometry data with the GNPS web platform. **a**) Two methods exist for the generation of molecular networks on the GNPS web platform: classical MN and feature-based molecular networking (FBMN). For both methods, mass spectrometry data files have to be converted to the mzML format using tools such as Proteowizard MSConvert^25^. The classical MN method runs entirely on the GNPS platform. In that method, MS^2^ spectra are clustered with MS-Cluster and the consensus MS^2^ spectra obtained are used for classical MN generation. In the case of FBMN, a feature detection and alignment tool is used to first process the LC-MS^2^ data (such as MZmine, MS-DIAL, XCMS, OpenMS, Progenesis QI, or MetaboScape) instead of using MS-Cluster (classical MN). Results are then exported and uploaded to the GNPS web platform for molecular networking analysis. **b**) Graphs showing the number of molecular networking jobs performed on GNPS. The upper graph shows the number of classical MN and FBMN jobs since 2016. The lower graph shows the number of FBMN jobs since its introduction in 2017 and key events accelerating its use.

**Fig. 2:**
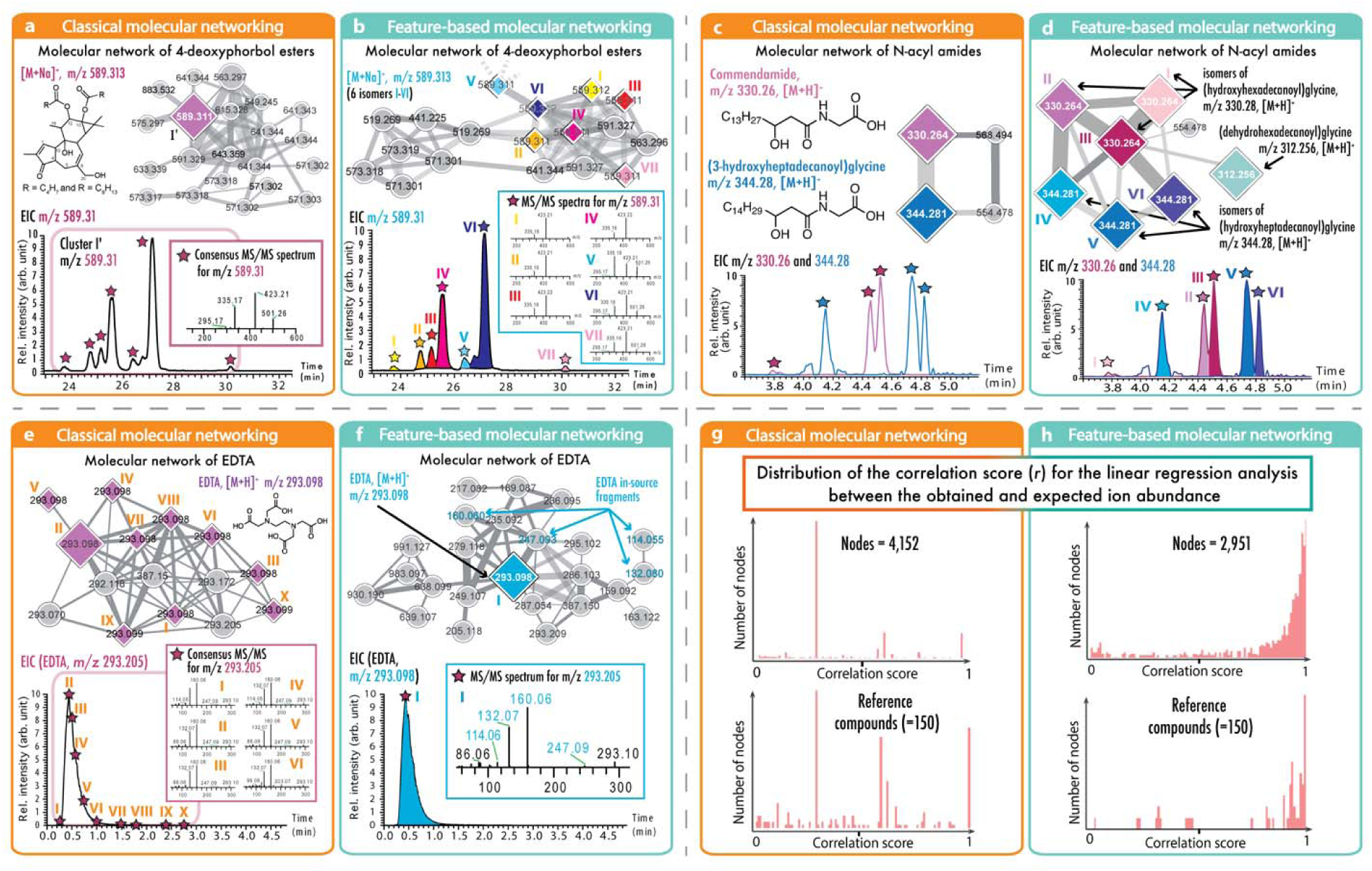
Comparisons of classical MN and feature-based molecular networking. In these examples, the node size corresponds to the spectral count in classical MN (orange boxes, left) or to the sum of LC-MS peak area in FBMN (blue boxes, right). Panel (**a**) displays the results from classical MN with the LC-MS^2^ data of *Euphorbia dendroides* plant samples; classical MN resulted in one node for the ion at *m/z* 589.313, while (**b**) FBMN performed after MZmine processing was able to detect seven isomers of this ion. Classical MN in the data of a cohort from the American Gut Project (**c**) showed two different *N*-acyl amides while the use of FBMN (**d**) processed with MZmine allowed the annotation of three different isomers per *N*-acyl amides. Classical MN (**e**) and FBMN (**f**) performed with OpenMS, were used to analyse the network of EDTA in plasma (373 samples) by compressing MS^2^ spectra of EDTA eluting over 2.5 min into one best-quality MS^2^ spectrum. FBMN recovered the molecular similarity of in-source fragments observed for EDTA, which were not displayed with classical MN, due to the fixed top-K rank for connected nodes (typically set to 10) of MS^2^ spectral similarity. Evaluation of quantitative performance using a serial dilution of serum reference sample (NIST1950SRM) analyzed with (**g**) classical MN and (**h**) FBMN. The plots are showing the correlation score (*r*) distribution between the observed and expected relative ion abundance. The upper charts present the distribution of the correlation score for all the nodes (features) generated, and the bottom charts show the distribution for 150 reference compounds. While classical MN uses the spectral count or the sum precursor ion count to estimate the molecular network node abundance, FBMN uses the LC-MS feature abundance (peak area or peak height), resulting in a more accurate estimation of the relative ion intensity.

By incorporating chromatographic information, FBMN reduced the complexity of molecular networking analysis. In non-targeted LC-MS^2^ data acquisition, the same precursor ion can be fragmented multiple times during chromatographic elution, which ultimately leads to multiple nodes representing the same compound in classical MN. Additionally, coeluting isobaric ions are often isolated and fragmented together leading to the generation of chimeric spectra which result in different MS^2^ spectra for one precursor ion. With FBMN, the integration of feature detection with the alignment of the mass spectrometry signal discerns that these different MS^2^ spectra are related to the same precursor molecule and selects a singular representative consensus spectrum for the feature. The benefit of using FBMN in such a case can be illustrated with the metal chelating agent ethylenediaminetetraacetic acid (EDTA) observed in the LC-MS^2^ analysis of plasma samples (Fig 2e), in which it is used as an anticoagulant agent. Classical MN resulted in 13 duplicated nodes with identical precursor *m/z values* in one molecular family, ten of which have spectral library matches to EDTA reference MS^2^ data (Fig. 2e and f). On the contrary, FBMN displays a unique representative MS^2^ spectrum that matches EDTA spectra in the library, because the multiple MS^2^ spectra detected were part of a single feature in the chromatographic dimension. The reduction of redundancy within the resulting molecular network simplifies the discovery of structurally related compounds.

While classical MN uses the spectral count or the sum precursor ion count, FBMN uses the LC-MS feature abundance (peak area or peak height), resulting in a more accurate estimation of the relative ion intensity. The method of FBMN simplifies, organizes the data, and adds relative quantitative information and precursor isotope patterns. FBMN enables robust statistical analysis by providing relative ion intensities across a dataset. This capacity is demonstrated with a serial dilution series dataset of the NIST1950 serum reference standard^13^, containing 150 spiked standards. Here, the LC-MS^2^ were processed with MZmine^14^ or OpenMS^15^ for FBMN (Fig 2g-h). A linear regression analysis was used to evaluate the relative quantification between classical MN and FBMN. Figure 2h shows that for FBMN, relative quantification has a correlation coefficient (*r*) value distribution mostly above 0.7, while this was not the case when the precursor ion abundance was obtained from classical MN via spectral counts (Fig 2g). The improved distribution of correlation coefficients towards 1 indicates a more linear response between concentration and ion abundance, which improves the accuracy and precision of quantification results. FBMN facilitates the direct application of existing statistical, visualization, and annotation tools, such as QIIME2^16^, MetaboAnalyst^17^, ili^18^, NAP^19^, MS2LDA^20^, MolNetEnhancer ^21^, and SIRIUS^22^.

FBMN further enables the creation of molecular networks from ion mobility spectrometry experiments coupled with LC-MS^2^ analysis. As an orthogonal separation method, the use of ion mobility offers additional resolving power to differentiate isomeric ions in the molecular network. The integration of ion mobility with FBMN on GNPS can currently be performed with MetaboScape, MS-DIAL^23^, and Progenesis QI. An example of such isomer separation using trapped ion mobility spectrometry (TIMS) coupled to LC-MS^2^ is shown in Supplementary Fig. 1.

Available on the GNPS web platform at https://gnps.ucsd.edu, FBMN is ideally suited for advanced molecular networking analysis, enabling the characterization of isomers, the incorporation of relative quantification, and the integration of ion mobility data. FBMN is the recommended way to analyse a single LC-MS^2^ metabolomics study, but care must be taken when applied across multiple studies due to different experimental conditions and possible batch effects. Moreover, the use of FBMN for the analysis of very large datasets (containing several thousand samples) is limited by the scalability of most feature detection and alignment software tools. Thus, while FBMN offers an improvement upon many aspects of molecular networking analysis, classical MN remains essential for repository-scale meta-analysis large dataset processing, and is convenient for rapid analysis of LC-MS^2^ data with less user defined parameters: one important aspect of molecular networks obtained with FBMN is the use of adequate processing steps and parameters, which otherwise could negatively impact the resulting molecular networks. To facilitate dissemination, education of the FBMN method, and the supported processing software, we created detailed tutorials and step-by-step instructions, available at https://ccms-ucsd.github.io/GNPSDocumentation/featurebasedmolecularnetworking.

The FBMN workflow offers not only automated spectral library search and spectral entry curation, but is also integrated with other annotation tools available on GNPS environment, such as MASST^24^, while promoting data analysis reproducibility by saving the FBMN jobs on the user’s private online workspace. The GNPS environment conveniently enables the user to evaluate different parameters and enables the sharing of the results via a web URL for publication.

## Methods

### Development of Feature-based Molecular Networking (FBMN)

The FBMN method consists of two main steps: 1) LC-MS feature detection and alignment, then 2) a dedicated molecular networking workflow on GNPS. Our first prototype for FBMN was developed with the Optimus workflow^12,18^ that uses OpenMS tools^15^ and the KNIME Analytics^26^ platform. Following step 1 (feature detection and alignment), two files are exported: a *feature quantification table* (.TXT format) and a *MS*^*2*^ *spectral summary* (.MGF format). The feature quantification table contains information about LC-MS features across all considered samples including a unique identifier (feature ID) for each feature, *m*/*z* value, retention time, and intensity. The *MS*^*2*^ *spectral summary* contains a list of MS^2^ spectra, with one representative MS^2^ spectrum per feature. The mapping information between the feature quantification table and the *MS*^*2*^ *spectral summary* is stored in these files using the feature ID and scan number, respectively. This simple mapping enables to relate LC-MS feature information or statistically derived results to the molecular network nodes. This approach was also used for the integration of other tools with FBMN, and does not require third party software like it was proposed in the past^27,28^. Finally, the FBMN workflow also supports the mzTab-M format^8^, a standardized output format designed for the report of metabolomics MS-data processing results. In this case, the mzTab-M file is used instead of *feature quantification table* and requires the input of the mzML files instead of the *MS*^*2*^ *spectral summary* file. Support for the mzTab-M format enables the possibility to perform FBMN with any existing and future processing tools that support this standardized format.

The FBMN workflow has been integrated into the GNPS ecosystem and thus benefits from the connection with other GNPS features, e.g. the possibility to perform automatic MS^2^ spectral library search, the direct addition and curation of library entries, the search of a spectrum against public datasets with MASST^24^, and the visualization of molecular networks directly in the web browser^29^ or with Cytoscape^30^. The FBMN workflow is available on the GNPS platform (https://gnps.ucsd.edu/) via a web interface (See Supplementary Fig. 2). Jobs are computed and stored on the computational infrastructure of the Center for Computational Mass Spectrometry at the University of California San Diego. Each finished job is saved in the private user space for future examination and has a permanent static link that enables data sharing and collaborative analyses. We strongly recommend the sharing of this static link along with publications using GNPS workflows to facilitate results accessibility and data analysis reproducibility. Instructions to perform FBMN with the supported tools are provided in the GNPS documentation (https://ccms-ucsd.github.io/GNPSDocumentation/featurebasedmolecularnetworking and Supplementary Fig. 3).

### FBMN with MZmine

MZmine^14^ is a popular open-source cross-platform software for mass spectrometry data processing with an advanced Graphical User Interface (GUI) that enables the users to visually optimize parameters and examine the results of each processing step. Moreover, MZmine allows for the export of a batch file containing all the steps and parameters used in the processing, thus enabling its reproducibility. To support FBMN in MZmine, the feature detection step (peak “Deconvolution module”) was modified to provide the ability to pair a feature with its MS^2^ scans using an *m/z* and retention time range defined by the user (Supplementary Fig. 4). Due to a new data structure and to support older projects (created with release < 2.38), an additional specific filtering module (*Group MS*^*2*^ *scans with features*) was developed to assign all MS^2^ scans to the features of existing peak list (see this video for instructions: https://www.youtube.com/watch?v=EL5pmFvpTFE). Moreover, a GNPS export and direct submission module was created (Supplementary Fig. 5) which offers two modes: 1) Export of the *feature quantification table* and the *MS*^*2*^ *spectral summary* file and 2) Direct FBMN analysis on the GNPS web platform (release 2.37+). The direct GNPS job submission generates all the files and uploads them together with an optional metadata table and default parameters (Supplementary Fig. 6) to the FBMN workflow on GNPS. By providing the user’s GNPS login credentials (optional), a new job can be created in the personal user space (https://www.youtube.com/watch?v=vFcGG7T_44E&list=PL4L2Xw5k8ITzd9hx5XIP94vFPxj1sSafB&index=4&t=0s). Otherwise, the user can be notified by email or directly redirected to the job webpage after the submission. With the option “*most intense*”, the GNPS Export uses the most intense MS^2^ spectrum as a representative spectrum for each LC-MS^2^ feature. When using the “merge MS/MS” spectra option (release 2.40+), a representative high quality MS^2^ spectrum is instead generated from all spectra and exported as a representative spectrum (Supplementary Note 1). The detailed documentation is available at https://ccms-ucsd.github.io/GNPSDocumentation/featurebasedmolecularnetworking-with-mzmine2/.

### FBMN with OpenMS

OpenMS is an open-source cross-platform software specifically designed for the flexible and reproducible analysis of high-throughput MS data analysis, including more than 200 tools for common mass spectrometric data processing tasks^15^. Building on our experience with the Optimus development, the integration of OpenMS and FBMN was achieved by creating a *GNPSExport* tool (TOPP tool) as a part of the OpenMS tool collection (https://github.com/Bioinformatic-squad-DorresteinLab/OpenMS). A detailed description of the *GNPSExport* module and how to use it for FBMN is available at the following webpage https://ccms-ucsd.github.io/GNPSDocumentation/featurebasedmolecularnetworking-with-openms/. In brief, after running an OpenMS non-targeted metabolomics pipeline, the *GNPSExport* TOPP tool can be applied to the consensusXML file resulting from *FeatureLinkerUnlabeledKD* or *FeatureLinkerUnlabeledQT* tools (alignment step), and the corresponding mzML files. For each consensusElement (LC-MS^2^ feature) in the consensusXML file, the *GNPSExport* generates one representative consensus MS^2^ spectrum that will be exported in the *MS*^*2*^ *spectral summary* file (using either the option “most intense” or “merged spectra”, see Supplementary Note 1). The *TextExport* tool is applied to the same consensusXML file to generate the *feature quantification table*. Note that the *GNPSExport* requires the use of the *IDMapper* tool on the featureXML files (from the feature detection step) prior to feature linking, in order to associate MS^2^ scans [peptide annotation in OpenMS terminology] with each feature. These MS^2^ scans are used by the GNPSExport for the generation of the representative MS^2^ spectrum. Additionally, the FileFilter has to be run on the consensusXML file, prior to the *GNPSExport*, in order to remove consensus Elements without associated MS^2^ scans. The two files exported (*feature quantification table* and *MS*^*2*^ *spectral summary*) can be directly used for FBMN analysis on GNPS. The OpenMS-GNPS workflow for metabolomics data processing was implemented as a python wrapper around OpenMS TOPP tools (https://github.com/Bioinformatic-squad-DorresteinLab/openms-gnps-tools), and released as a workflow (https://github.com/Bioinformatic-squad-DorresteinLab/openms-gnps-workflow) on the GNPS/MassIVE web platform^31^. OpenMS version 2.4.0 was used^15^. The OpenMS + GNPS workflow can be accessed and run here: https://proteomics2.ucsd.edu/ProteoSAFe/.

### FBMN with XCMS

The XCMS package (https://github.com/sneumann/xcms for the most recent version) is one of the most widely used software for processing of mass spectrometry-based metabolomics data^32^. The integration of XCMS and FBMN is currently possible using a custom utility function “formatSpectraForGNPS” creating the *MS*^*2*^ *spectral summary.* This function is available on the following GitHub repository https://github.com/jorainer/xcms-gnps-tools and is compatible with the CAMERA algorithm for isotopes and adduct annotation^33^. Representative XCMS R scripts in markdown and Jupyter notebook formats are available in the following GitHub repository https://github.com/DorresteinLaboratory/XCMS3_FeatureBasedMN. The two exported files (*feature quantification table* and *MS*^*2*^ *spectral summary*) can be directly used for FBMN analysis on GNPS. The detailed documentation is available at https://ccms-ucsd.github.io/GNPSDocumentation/featurebasedmolecularnetworking-with-xcms3/.

### FBMN with MS-DIAL

MS-DIAL is an open-source mass spectrometry data processing software^23^ (available for Windows only, http://prime.psc.riken.jp/Metabolomics_Software/MS-DIAL/). The integration of MS-DIAL and FBMN was made possible since ver. 2.68 by exporting the “Alignment results” using the “GNPS export” option. In addition to LC-MS^2^ data processing, MS-DIAL can process data from SWATH-MS^2^ (data-independent LC-MS^2^ acquisition), and ion mobility spectrometry coupled to LC-MS^2^. The two files exported (*feature quantification table* and *MS*^*2*^ *spectral summary*) can be directly used for FBMN analysis on GNPS. A video tutorial on the use of MS-DIAL for FBMN is available at https://www.youtube.com/watch?v=hxk40jwAkcc&t=7s. The detailed documentation is available at https://ccms-ucsd.github.io/GNPSDocumentation/featurebasedmolecularnetworking-with-ms-dial/.

### FBMN with MetaboScape

MetaboScape is a proprietary mass spectrometry metabolomics data processing software commercialized by Bruker and available on Windows. MetaboScape can perform feature detection, alignment and annotation of non-targeted LC-MS^2^ data acquired on Bruker mass spectrometers. Support for the processing of trapped ion mobility spectrometry (TIMS) coupled to non-targeted LC-MS^2^ (LC-TIMS-MS^2^) was added in MetaboScape 4.0, which results in LC-TIMS-MS features. Feature-based molecular networking can be performed on LC-MS^2^ or LC-TIMS-MS^2^ data by exporting the *feature quantification table* and *MS*^*2*^ *spectral summary* from the “bucket table” using the “Export to GNPS format” function. These files can be uploaded to GNPS for FBMN analysis. Information from MetaboScape, such as the Collision Cross Section values, or other spectral annotations can be mapped into the molecular networks using Cytoscape^30^. The detailed documentation is available at https://ccms-ucsd.github.io/GNPSDocumentation/featurebasedmolecularnetworking-with-metaboscape/

### FBMN with Progenesis QI

Progenesis QI is a proprietary feature detection and alignment software developed by Nonlinear Dynamics (Waters) that is compatible with various proprietary and open mass spectrometry data formats. Progenesis QI can perform feature detection, alignment and annotation of non-targeted LC-MS^2^ data acquired either in data-dependent acquisition (DDA) or data independent analysis (DIA, such as MS^E^), and can also utilize the ion mobility spectrometry (IMS) dimension. FBMN can be performed on any of these data types processed with Progenesis QI (ver 4.0), by exporting the *feature quantification table* (.CSV format) and the *MS*^*2*^ *spectral summary* (.MSP format). These two files can be exported from the “Identify Compounds” submenu by using the function “Export compound measurement” and “Export fragment database”, respectively. These files can be uploaded to GNPS for FBMN analysis. Information from Progenesis QI, such as the Collision Cross Section values, or other spectral annotations can be mapped into the molecular networks using Cytoscape^30^. The detailed documentation is available at https://ccms-ucsd.github.io/GNPSDocumentation/featurebasedmolecularnetworking-with-progenesisQI/.

### FBMN makes it possible to resolve isomers in a drug lead discovery effort

The examination of the LC-MS^2^ data (<u>MSV000080502</u>) from the *Euphorbia dendroides* plant extract showed the presence of numerous chromatographic peaks for ions in the range *m/z* 500-900, corresponding to diterpene ester derivatives. These specialized metabolites consist of a polyhydroxylated diterpene core acylated with various acidic moieties, that are typically found as positional isomers based on their acylation pattern^34^. The extracted ion chromatogram (EIC) for the ion *m*/*z* 589.31 in the *Euphorbia dendroides* extract data (Supplementary Fig. 7) shows the presence of at least seven distinct LC-MS peaks between 24.5 and 27.3 min, including five peaks with an associated MS^2^ spectra. The analysis of the extract and the fractions where these molecules were originally isolated (fractions 13 and 14) with <u>classical MN</u> resulted in a <u>molecular network</u> with two nodes for the *m/z* 589.31 ions (Fig. 2a and Supplementary Fig. 8). These MS^2^ spectra (cluster index <u>5352</u> and <u>5354</u>) resulted from merging 96 fragmentation spectra spanning from 23.6 to 26.5 min by MS-Cluster (Fig. 2b and Supplementary Fig. 9). Close examination of the clustered spectra revealed that while all MS^2^ spectra for the precursor *m/z* 589.31 present fragment ions *m/z* 501.26, 423.21, 335.16, and 295.17, three distinct spectral types could be established based on the ions relative intensities (Supplementary Fig. 10). FBMN of the dataset with MZmine processing (<u>see the GNPS job</u>) enabled the differentiation of the MS^2^ spectra of seven isomers (Figure 2b and Supplementary Fig. 11 for the <u>molecular network view</u>). A detailed discussion on the differences observed between the two methods can be found in the Supporting Information (Supplementary Note 2 and Supplementary Table 1). Interestingly, in the original study^12^ OpenMS was used for FBMN and resulted in the observation of three different positional isomers instead of seven, which shows that different processing methods can lead to different results with FBMN. These three isomers were subsequently isolated and differed by the position of one double bond on the C-12 acyl chain, or from carbon C-4 configuration^12^. Because FBMN connects the accurate relative abundance of the ions across the fractions and the molecular networks, it allowed to create bioactivity-based molecular networks^12^, which were used to predict and target potentially antiviral compounds. For detailed description of the extraction, mass spectrometry analysis, and structural elucidation, see the original manuscript^12^. The MZmine project and parameters used can be accessed on the MassIVE submission (<u>MSV000080502</u>).

### FBMN resolves isomers in large scale metabolomics studies

FBMN was applied on a cohort of the American Gut Project (AGP), a citizen-scientist research project that enabled the observation of the commendamide in humans, along with other new *N*-acyl amide derivatives using molecular networking^11^. Commendamide is a recently discovered bacterial *N*-acyl amide that was shown to modulate host metabolism via G-protein-coupled receptors (GPCRs) in the murine intestinal tract^35^.

The use of FBMN for the AGP data (Figure 2d) made possible to observe the presence of two additional commendamide isomers (*m*/*z* 330.26) and of an analogue (hydroxyheptadecanoyl)glycline (*m*/*z* 344.28), while classical MN resulted in the observation of one single consensus spectrum for each compound (Figure 2c). In addition, FBMN allowed to observe a putative commendamide derivative (dehydrohexadecanoyl)glycine (<u>CCMSLIB00005436498</u> and Supplementary Fig. 12) in the commendamide molecular network. The sample collection and mass spectrometry acquisition methods are described in the original manuscript^11^. The data were downloaded from MassIVE (<u>MSV000080186</u>) and processed with MZmine (2.37). The MZmine project along with parameters and export files were deposited to the MassIVE repository (<u>MSV000084095</u>). The chromatograms for *m*/*z* 330.26 and *m*/*z* 344.28 displayed in Figure 2c-d are from samples 43076_P3_RB9_01_314.mzML and 38131_P5_RA4_01_538.mzML, respectively. Chromatograms were exported with MZmine. The results were exported with the “Export for/Submit to GNPS” module for FBMN analysis on GNPS. The corresponding job can be accessed here: https://gnps.ucsd.edu/ProteoSAFe/status.jsp?task=0a8432b5891a48d7ad8459ba4a89969f (only logged users can see all the input files). The mzML files were used for the classical MN job can be accessed here: https://gnps.ucsd.edu/ProteoSAFe/status.jsp?task=3c27e43d908c4044bace405cc394cd25.

### FBMN reduces spectral redundancy and deobfuscates spectral similarity relationships: the case of EDTA

The benefit of using FBMN can be illustrated with the metal chelating agent ethylenediaminetetraacetic acid (EDTA), widely used in beauty products, food, and scientific protocols. A search for its occurrences in public spectral datasets with the mass spectrometry search tool (MASST)^24^ showed that it <u>is frequently observed</u> in plasma samples where it is used during the sample preparation. We took one of the public human serum sample datasets (<u>MSV00008263</u>) where EDTA was observed. For a detailed description of the protocol and mass spectrometry parameters, see Supplementary Note 3. The analysis of the data with classical MN showed that the EDTA ions are found in two molecular networks. One network consists of <u>[M+H]</u>^+^ spectra and the other of <u>[M+Na]</u>^+^ spectra. Interestingly, each of these networks have one node with a large number of clustered spectra (node 91205 for 4655 spectra, and node 116470 for 571 spectra, respectively), but yet EDTA ions are represented by multiple nodes although these nodes have the same precursor ion mass and retention time. Detailed analysis showed that while the median pairwise cosine values between EDTA spectra are high (median value of 0.93 and 0.94), the spectra are not clustering into a single node. Examination of the multiple fragmentation spectra for EDTA ions showed that some 1) are chimeric spectra “contaminated” by fragment ions produced by co-eluting isobaric ions, and 2) that other spectra were dominated by low intensity fragment ions resulting from MS^2^ spectra acquired at low intensity. The method of FBMN was applied on that same dataset using the OpenMS-GNPS workflow (<u>see the job</u>), and the results showed that it efficiently reduces the appearance of these redundant node patterns from the same molecule (<u>see the FBMN job</u>, Figure 2f), both for the molecular networks containing the [<u>M+H</u>]^+^ and [<u>M+Na</u>]^+^ spectra. FBMN recovers the molecular similarity of in-source fragments observed for EDTA, which were not displayed with classical MN, as they now fall within the top-K rank (typically set to 10) of MS^2^ spectral similarity considered in the network topology. The parameters used for OpenMS tools can be accessed in the OpenMS-GNPS job (<u>see the job</u>). OpenMS ver. 2.4.0 was used^15^.

### FBMN enables the use of relative quantification in the molecular networks

While classical MN uses the spectral count or the sum of precursor ion intensity to estimate the ion abundance, FBMN uses the accurate ion intensities obtained from LC-MS feature detection. The FBMN method brings in ion abundance across all samples by using the value of the chromatographic peak area or peak height as determined by the LC-MS feature detection and alignment software. Using a serial dilution of the NIST 1950 serum reference metabolome sample^13^ analyzed on an Orbitrap mass spectrometer (Q Exactive, ThermoFisher) and processed with OpenMS or MZmine, we show the linearity of the relative quantification with FBMN and the improvement compared to classical MN (Figure 2h). The sample preparation and mass spectrometry methods are described in Supplementary Note 4. The files along with the parameters for MZmine are available on the following MassIVE repository (<u>MSV000084092</u>). The linear regression analysis was performed with python 2 (ver. 2.7.15) with the *LinearRegression* function of the *sklearn* package (ver. 0.20.1)^36^. The parameters used for OpenMS can be obtained from the following job link: https://proteomics2.ucsd.edu/ProteoSAFe/status.jsp?task=aae71e9b72cf431d9b2606170c3f7a7d. The molecular networking jobs can be accessed here: classical MN (https://gnps.ucsd.edu/ProteoSAFe/status.jsp?task=daf3f0d7cec94104b2c9001739964c31), MZmine processing (https://gnps.ucsd.edu/ProteoSAFe/status.jsp?task=f443cad083be4979aedd2af0f97b9fe9) and OpenMS processing (https://gnps.ucsd.edu/ProteoSAFe/status.jsp?task=2c48a477ec094123987cdf90db4be8e4).

### FBMN enables molecular networking with ion mobility spectrometry

The sample NIST 1950 serum^13^ was analyzed using a timsTOF Pro (Bruker Daltonics, Bremen) in data-dependent acquisition mode using PASEF (Parallel Accumulation-Serial Fragmentation)^37^. The data were then processed with MetaboScape (ver. 5.0) and the results were exported for FBMN analysis on GNPS. The mass spectrometry acquisition method, data, and parameters used for the processing were deposited on MassIVE (<u>MSV000084402</u>). Classical MN were annotated with the GNPS^6^, NIST17 and LipidBlast^38^ spectral libraries https://gnps.ucsd.edu/ProteoSAFe/status.jsp?task=f2adc2cf33c646548798d0e285197a96). Lipid annotation in MetaboScape was performed using SimLipid (ver. 6.04, Premier Biosoft, Palo Alto) and mapped to the FBMN (https://gnps.ucsd.edu/ProteoSAFe/status.jsp?task=0d89db67b0974939a91cb7d5bfe87072). The molecular networks were visualized with Cytoscape ver. 3.7.1^30^, and the results are presented in the Supplementary Information (Fig. S1). Classical MN were annotated with the GNPS^6^, NIST17 and LipidBlast^38^ spectral libraries https://gnps.ucsd.edu/ProteoSAFe/status.jsp?task=f2adc2cf33c646548798d0e285197a96). FBMN were annotated by mapping annotations from the MetaboScape annotation engine (https://gnps.ucsd.edu/ProteoSAFe/status.jsp?task=0d89db67b0974939a91cb7d5bfe87072).

### Integration with other computational mass spectrometry annotation tools

The MGF file format is accepted by numerous computational mass spectrometry annotation tools. The use of these tools with the MS^2^ spectral summary file enables subsequent direct mapping of these annotations to the molecular networks produced by the feature-based molecular networking method. These tools include SIRIUS^22^, DEREPLICATOR,^39,40^ NAP,^19^ MS2LDA^20^, MolNetEnhancer^21^ (see Supplementary Note 5), as well as other software such as MetWork^41^, CFM-ID^42^, MetGem^43^, MetFrag^44^.

### Large dataset processing with OpenMS and XCMS

The processing of large metabolomics datasets (more than a thousand samples) is limited by the scalability of existing LC-MS feature detection tools, especially those based on a GUI (such as MZmine and MS-DIAL). We showed that with specific peak picking parameters the use of XCMS or OpenMS enables using FBMN for large metabolomics study (<u>MSV000080030</u>, approximately 2,000 samples). See the Supplementary Note 6, Supplementary Table 2, and Supplementary Fig. 13.

## Code availability

The FBMN workflow is available as web-interface on the GNPS web platform (https://gnps-quickstart.ucsd.edu/featurebasednetworking). The workflow code is open source and available on GitHub (https://github.com/CCMS-UCSD/GNPS_Workflows/tree/master/feature-based-molecular-networking). It is released under the licence of The Regents of the University of California and free for non-profit research (https://github.com/CCMS-UCSD/GNPS_Workflows/blob/master/LICENSE). The workflow was written in Python (ver. 3.7) and deployed with the ProteoSAFE workflow manager employed by GNPS (http://proteomics.ucsd.edu/Software/ProteoSAFe/). We also provide documentation, support, example files, and additional information on the GNPS documentation website (https://ccms-ucsd.github.io/GNPSDocumentation/featurebasedmolecularnetworking/). The source code of the GNPSExport module in MZmine is available at (https://github.com/mzmine/mzmine2) under the GNU General Public License. The source code of the GNPSExport tool in OpenMS is available at (https://github.com/Bioinformatic-squad-DorresteinLab/OpenMS) under the BSD licence. The source code for the GNPSExport custom function for XCMS is available at https://github.com/jorainer/xcms-gnps-tools under the GNU General Public License.

## Data availability

The LC-MS^2^ data for the *Euphorbia dendroides* dataset, along with the MZmine project and parameters used can be accessed on the MassIVE submission (<u>MSV000080502</u>, Creative Commons CC0 1.0 Universal license). The classical MN and FBMN jobs can be accessed via the GNPS website at https://gnps.ucsd.edu/ProteoSAFe/status.jsp?task=189e8bf16af145758b0a900f1c44ff4a and https://gnps.ucsd.edu/ProteoSAFe/status.jsp?task=672d0a5372384cff8c47297c2048d789, respectively. The LC-MS^2^ data for the American Gut Project (AGP) were downloaded from MassIVE (<u>MSV000080186</u> Creative Commons CC0 1.0 Universal license) and processed with MZmine (2.37). The MZmine project along with parameters and export files were deposited (<u>MSV000084095</u>, Creative Commons CC0 1.0 Universal license). The classical MN and FBMN jobs can be accessed at https://gnps.ucsd.edu/ProteoSAFe/status.jsp?task=3c27e43d908c4044bace405cc394cd25 and https://gnps.ucsd.edu/ProteoSAFe/status.jsp?task=0a8432b5891a48d7ad8459ba4a89969f, respectively. The LC-MS^2^ data for the EDTA case are available on the MassIVE submission (<u>MSV00008263</u>, Creative Commons CC0 1.0 Universal license). The classical MN job can be accessed at https://gnps.ucsd.edu/ProteoSAFe/status.jsp?task=fbac1a5061ba4ad683a284ef55d45df6). The OpenMS and the FBMN job at https://proteomics2.ucsd.edu/ProteoSAFe/status.jsp?task=83a0a417a49b4b76b61e9a8191a6ea2d at https://gnps.ucsd.edu/ProteoSAFe/status.jsp?task=8f40420c11694cf9ab06fdf7a5a4c53b, respectively. The mass spectrometry acquisition method, data, and parameters used for the processing of the serum analysis with the timsTOF mass spectrometer were deposited (<u>MSV000084402</u>). Classical MN and FBMN jobs can be accessed here: https://gnps.ucsd.edu/ProteoSAFe/status.jsp?task=f2adc2cf33c646548798d0e285197a96, and https://gnps.ucsd.edu/ProteoSAFe/status.jsp?task=0d89db67b0974939a91cb7d5bfe87072, respectively.

## Supporting information

Supplementary informations

## Author Contributions

L.F.N., D.P., M.W., and P.D. conceived the method and supervised its implementation and wrote the manuscript.

L.F.N and D.P contributed equally to the work.

I.P., L.F.N., M.E., and T.A. created the FBMN prototype in Optimus. M.W., L.F.N. D.P. and Z.Z. created the FBMN workflow on GNPS.

R.S., L.F.N, M.W., D.P., A.K., M.F, Z.Z., A.S., and T.P. developed the GNPS Export module in MZmine.

K.D., A.K., M.L., and S.B. developed the spectral clustering algorithm and SIRIUS export in MZmine.

A.S., and L.F.N. created the GNPS Export tool in OpenMS, with the guidance from F.A, O.A., and O.K.

J.R. and M.Wit. created the XCMS export tool.

H.T, M.W. and L.F.N. made possible the integration with MS-DIAL.

L.F.N., A.B., H.N., F.Z. and T.D. made possible the integration with MetaboScape.

M.W., G.I., B.S., S.W.M. and J.M. made possible the integration with Progenesis QI.

F.V. performed the mass spectrometry for the plasma and NIST1950SRM samples.

A.A. performed the mass spectrometry for the American Gut Project samples.

A.K.J, L.F.N, and A.Tri. analyzed the results of the plasma samples.

J.R and L.F.N. performed the XCMS processing of the forensic dataset.

L.F.N. and M.W. created the general documentation.

D.P., L.F.N. and R.d.S. created the MZmine documentation.

K.B.K., H.Y. created the MS-DIAL documentation.

F.V., J.M.G. K.W., and A.K.J. prepared the MS-DIAL video tutorial.

M.W., R.S., D.P. prepared the MZmine video tutorials.

M.E., R.d.S., J.R., O.M., and S.N. created the XCMS documentation.

L.F.N and A.S. created the OpenMS documentation.

L.F.N, N.H.N., and T.D, created the MetaboScape documentation.

M.C., and L.-I. M. documented the FBMN interface workflow.

M.N.-E., I.K., and C.M created the Cytoscape documentation.

H.M, A.G., M.W. and L.F.N. made the integration with DEREPLICATOR.

M.W., J.J.J.v.d.H, M.E. and S.R. made the integration with MS2LDA.

R.d.S made the integration with NAP.

M.M., N.B., X.C., V.V.P., N.G. R.A.Q, A.A., Z.K., and S.N. tested and provided suggestions on how to improve the methods.

J.J.J.v.d.H., V.P., T.A., A.K.J., T.P., A.L.G, L.-I.M., P.-M.A., S.B., and S.N. improved the manuscript.

All authors have contributed to the final manuscript.

## Ethics/COI declaration

Pieter C. Dorrestein is a scientific advisor for Sirenas LLC.

Mingxun Wang is a consultant for Sirenas LLC and the founder of Ometa labs LLC. Tomáš Pluskal is a consultant for Ginkgo Bioworks.

Alexander Aksenov is a consultant for Ometa labs LLC.

Theodore Alexandrov is on the Scientific Advisory Board of SCiLS, a Bruker company.

Kai Dührkop, Marcus Ludwig, Markus Fleischauer and Sebastian Böcker are founders of Bright Giant GmbH.

Aiko Barsch, Sven W. Meyer, Heiko Neuweger and Florian Zubeil are employes of Bruker Daltonics GmbH.

Giorgis Isaac, Jonathan McSayles, and Bindesh Shrestha are employees of Waters Corporation.

## Acknowledgements

We gratefully acknowledge financial support by: the U.S. National Institutes of Health for the Center for Computational Mass Spectrometry grant (P41 GM103484), the reuse of metabolomics data (R03 CA211211), and the tools for rapid and accurate structure elucidation of natural products (R01 GM107550) to P.D; the European Union’s Horizon 2020 grants 704786 (MSCA-GF to L.F.N) 634402, 777222 (T.A. and I.P.) and the ERC Consolidator grant METACELL (T.A.). The German Research Foundation (DFG) with grant number PE 2600/1 to D.P.; S.N. acknowledges funding from Bundesministerium für Bildung und Forschung (FKZ 031L0107) and the European Commission (EC654241); NSF grant IOS-1656481 to PCD and AMCR; O.A., acknowledge the funding from: the Bundesministerium für Ernährung und Landwirtschaft (FKZ 2816501214), the Bundesministerium für Wirtschaft und Energie (FKZ AiF18475N), the Bundesministerium für Bildung und Forschung (FKZ 031A430C), and the European Commission (823839) that also benefited to F.A. and O.K.; S.B. acknowledges funding from Deutsche Forschungsgemeinschaft (BO 1910/20); J.J.J.v.d.H. was supported by an Accelerating Scientific Discoveries Grant funded by the Netherlands eScience Center [NLeSC] (No. ASDI.2017.030). Z.K. was supported by the project International Mobility of Researchers (CZ.02.2.69/0.0/0.0/16_027/0007990). The authors would like to thank Nils Hoffman for maintaining the mzTab-M format.

