## Supplementary informations for "Feature-based Molecular Networking in the GNPS Analysis Environment"

### Supplementary information

Louis Felix Nothias,[1,2,#] Daniel Petras,[1,2,3,#] Robin Schmid,[4] Kai Dührkop,[5] Johannes Rainer,[6] Abinesh Sarvepalli,[1,2] Ivan Protsyuk,[7] Madeleine Ernst,[1,2,8] Hiroshi Tsugawa,[9,10] Markus Fleischauer,[5] Fabian Aicheler,[11,12] Alexander Aksenov,[1,2] Oliver Alka,[11,12] Pierre-Marie Allard,[13] Aiko Barsch,[14] Xavier Cachet,[15] Mauricio Caraballo,[1,2] Ricardo R. Da Silva,[2,16] Tam Dang,[2,17] Neha Garg,[18] Julia M. Gauglitz,[1,2] Alexey Gurevich,[19] Giorgis Isaac,[20] Alan K. Jarmusch,[1,2] Zdeněk Kameník,[21] Kyo Bin Kang,[1,2,22] Nikolas Kessler,[14] Irina Koester,[1,2,3] Ansgar Korf,[4] Audrey Le Gouellec,[23] Marcus Ludwig,[5] Christian Martin H.,[24] Laura-Isobel McCall,[25] Jonathan McSayles,[26] Sven W. Meyer,[14] Hosein Mohimani,[27] Mustafa Morsy,[28] Oriane Moyne,[23,29] Steffen Neumann,[30,31] Heiko Neuweger,[14] Ngoc Hung Nguyen,[1,2] Melissa Nothias-Esposito,[1,2] Julien Paolini,[32] Vanessa V. Phelan,[33] Tomáš Pluskal,[34] Robert A. Quinn,[35] Simon Rogers,[36] Bindesh Shrestha,[19] Anupriya Tripathi,[1,29,37] Justin J.J. van der Hooft,[1,2,38] Fernando Vargas,[1,2] Kelly C. Weldon,[1,2,39] Michael Witting,[40] Heejung Yang,[41] Zheng Zhang,[1,2] Florian Zubeil,[14] Oliver Kohlbacher,[11,12,42,43] Sebastian Böcker,[5] Theodore Alexandrov,[1,2,7] Nuno Bandeira,[1,2,44] Mingxun Wang,[1,2,44]* and Pieter C. Dorrestein[1,2,29,39,]*

1. Skaggs of Pharmacy and Pharmaceutical Sciences, University of California San Diego, La Jolla, San Diego, CA, USA
2. Collaborative Mass Spectrometry Innovation Center, University of California San Diego, La Jolla, San Diego, CA, USA
3. Scripps Institution of Oceanography, University of California San Diego, La Jolla, CA, USA
4. Institute of Inorganic and Analytical Chemistry, University of Münster, Münster, Germany
5. Chair for Bioinformatics, Friedrich-Schiller-University Jena, Jena, Germany
6. Institute for Biomedicine, Eurac Research, Affiliated Institute of the University of Lübeck, Bolzano, Italy
7. Structural and Computational Biology Unit, European Molecular Biology Laboratory, Heidelberg, Germany
8. Center for Newborn Screening, Department of Congenital Disorders, Statens Serum Institut, Copenhagen, Denmark
9. RIKEN Center for Sustainable Resource Science, Yokohama, Kanagawa, Japan
10. RIKEN Center for Integrative Medical Sciences, Yokohama, Kanagawa, Japan
11. Applied Bioinformatics, Department of Computer Science, University of Tübingen, Tübingen, Germany
12. Institute for Translational Bioinformatics, University Hospital Tübingen, Tübingen, Germany
13. Department of Phytochemistry and Bioactive Natural Products, University of Geneva, Geneva, Switzerland
14. Bruker Daltonics, Bremen, Germany
15. Equipe PNAS, UMR 8038 CiTCoM CNRS, Faculté de Pharmacie de Paris, Université Paris Descartes, Paris, France
16. Department of Physics and Chemistry, School of Pharmaceutical Sciences of Ribeirão Preto, University of São Paulo, Ribeirão Preto, Brazil

17. Technische Universität Berlin, Faculty II Mathematics and Natural Sciences, Institute of Chemistry, Berlin, Germany
18. School of Chemistry and Biochemistry, Center for Microbial Dynamics and Infection, Georgia Institute of Technology, Atlanta, GA, USA
19. Center for Algorithmic Biotechnology, Institute of Translational Biomedicine, St. Petersburg State University, St. Petersburg, Russia
20. Waters Corporation, Milford, MA, USA
21. Institute of Microbiology of the Czech Academy of Sciences, Prague, Czech Republic
22. College of Pharmacy, Sookmyung Women's University, Seoul, Republic of Korea
23. Univ. Grenoble Alpes, CNRS, Grenoble INP, CHU Grenoble Alpes, TIMC-IMAG, Grenoble, France
24. Centro de Biodiversidad y Descubrimiento de Drogas, INDICASAT AIP, Panama, Republic of Panama
25. Department of Chemistry and Biochemistry, Department of Microbiology and Plant Biology and Laboratories of Molecular Anthropology and Microbiome Research, University of Oklahoma, USA
26. Nonlinear Dynamics, Milford, MA, USA
27. Computational Biology Department, School of Computer Sciences, Carnegie Mellon University, Pittsburgh, Pennsylvania, USA
28. Department of Biological and Environmental Sciences, University of West Alabama, Livingston, USA
29. Department of Pediatrics, University of California San Diego, La Jolla, San Diego, CA, USA
30. Bioinformatics and Scientific Data, Leibniz Institute of Plant Biochemistry, Halle, Germany
31. German Centre for Integrative Biodiversity Research (iDiv) Halle-Jena-Leipzig, Germany
32. Laboratoire de Chimie des Produits Naturels, UMR CNRS SPE, Université de Corse Pascal Paoli, France
33. Skaggs School of Pharmacy and Pharmaceutical Sciences, University of Colorado, Denver, Aurora, CO, USA
34. Whitehead Institute for Biomedical Research, Cambridge, MA, USA
35. Department of Biochemistry and Molecular Biology, Michigan State University, East Lansing, 48823, MI, USA
36. School of Computing Science, University of Glasgow, Glasgow G12 8QQ, UK
37. Division of Biological Sciences, University of California San Diego, La Jolla, CA, USA
38. Bioinformatics Group, Wageningen University, Wageningen, the Netherlands
39. Center for Microbiome Innovation, University of California, San Diego, La Jolla, CA, USA
40. Research Unit Analytical BioGeoChemistry, Helmholtz Zentrum München
41. College of Pharmacy, Kangwon National University, Republic of Korea
42. Institute for Bioinformatics and Medical Informatics, University of Tübingen, Tübingen, Germany
43. Biomolecular Interactions, Max Planck Institute for Developmental Biology, Tübingen, Germany
44. Department of Computer Science and Engineering, University of California San Diego, CA, USA

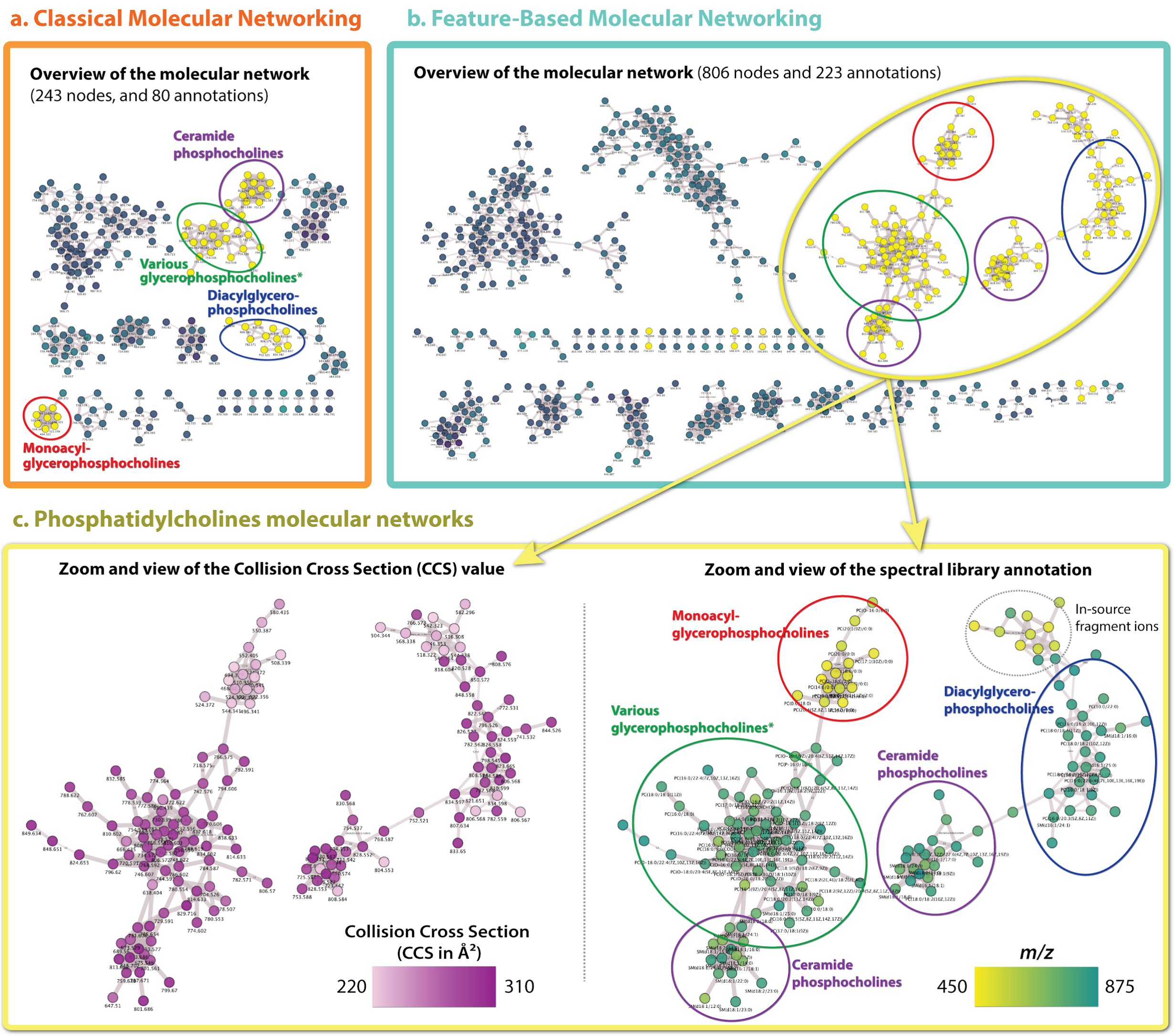


**Supplementary Figure 1.** Analysis of trapped ion mobility spectrometry (TIMS) data of the reference serum sample (NIST 1950 SRM) processed used with Feature-Based Molecular Networking (FBMN) on GNPS. The data were collected on a tims-TOF Pro (Bruker Daltonics, Bremen) in Parallel-Accumulation Serial Fragmentation (PASEF) mode, and processed with MetaboScape. (a) Molecular networks obtained with classical molecular networking; (b) molecular networks obtained with FBMN; and (c) views of the phosphatidylcholine molecular networks with FBMN (the node color gradient to the left shows the cross-collision section value, and the *m*/*z* value to the right). Results showed that the use of classical molecular networking drastically reduces the number of nodes (-70%) and spectral annotations (-65%) compared to FBMN. This can be explained by the fact that many isomers are present amongst the lipids annotated (mostly phosphatidylcholines). These isomers tend to produce similar $MS^2$ spectra which can be merged by MS-Cluster when performing classical molecular networking. FBMN ensures that the “LC-TIMS-MS” features detected by MetaboScape are preserved which enables the visualization of the Collision Cross Section value in the molecular networks.

**Supplementary Note 1: Methods for the generation of a representative MS$^2$ spectrum from a LC-MS$^2$ file**

The selection of the representative MS$^2$ spectrum in the *MS$^2$ spectral summary* file can be performed with different methods. Available in all tools supported, the most basic method named "Most intense" use the MS$^2$ spectrum with the highest precursor ion intensity or total ion current (in the specified *m*/*z* and retention time range) as the representative MS$^2$ spectrum. An experimental spectral "clustering" method for the creation of the representative MS$^2$ spectrum was implemented in MZmine, OpenMS, and XCMS. The clustering of MS$^2$ spectra is an intense research topics in proteomics[1], and more recently invested in metabolomics[2]. It has been shown that the clustering of MS$^2$ spectra can improve the identification rate in proteomics (done by spectral library matching), by increasing the signal/noise ratio of conserved fragment peaks[3]. However, clustering approaches can be negatively impacted by the presence of 1) low quality spectra, and 2) chimeric spectra. Low quality spectra are consisting of low signal/noise ratio fragment ions caused by poor fragmentation (producing fragment ions observed at detection limit of the instrument), or by fragmentation scans triggered at low intensity. Chimeric spectra are produced by the simultaneous selection and fragmentation of two isobaric co-eluting compounds, resulting in a mixed "chimeric" MS$^2$ spectrum. The spectral clustering method implemented in MZmine (Merge option in the GNPS/Sirius Export modules) and OpenMS (option "merge spectra", in the GNPSExport tool) works as follows: for each LC-MS/MS feature, the purity of each fragmentation spectra is calculated with a function inspired by msPurity[4]. In brief, adjacent MS$^1$ scans are examined to determine if other isobaric ions were co-fragmented. In these MS$^1$ scans, the ratio between the precursor ion intensity and the other isobaric ions in the ion isolation range is calculated. Then the purest MS$^2$ spectrum (highest purity score) is selected as the reference spectrum, and pairwise comparison (cosine score) is computed between the purest spectrum and all the other MS$^2$ spectra for the feature. All the MS$^2$ spectra reaching a cosine score threshold are then merged in one consensus or representative MS$^2$ spectrum. The mass accuracy, isolation width for the ion filtering and the cosine score, are defined by the user.

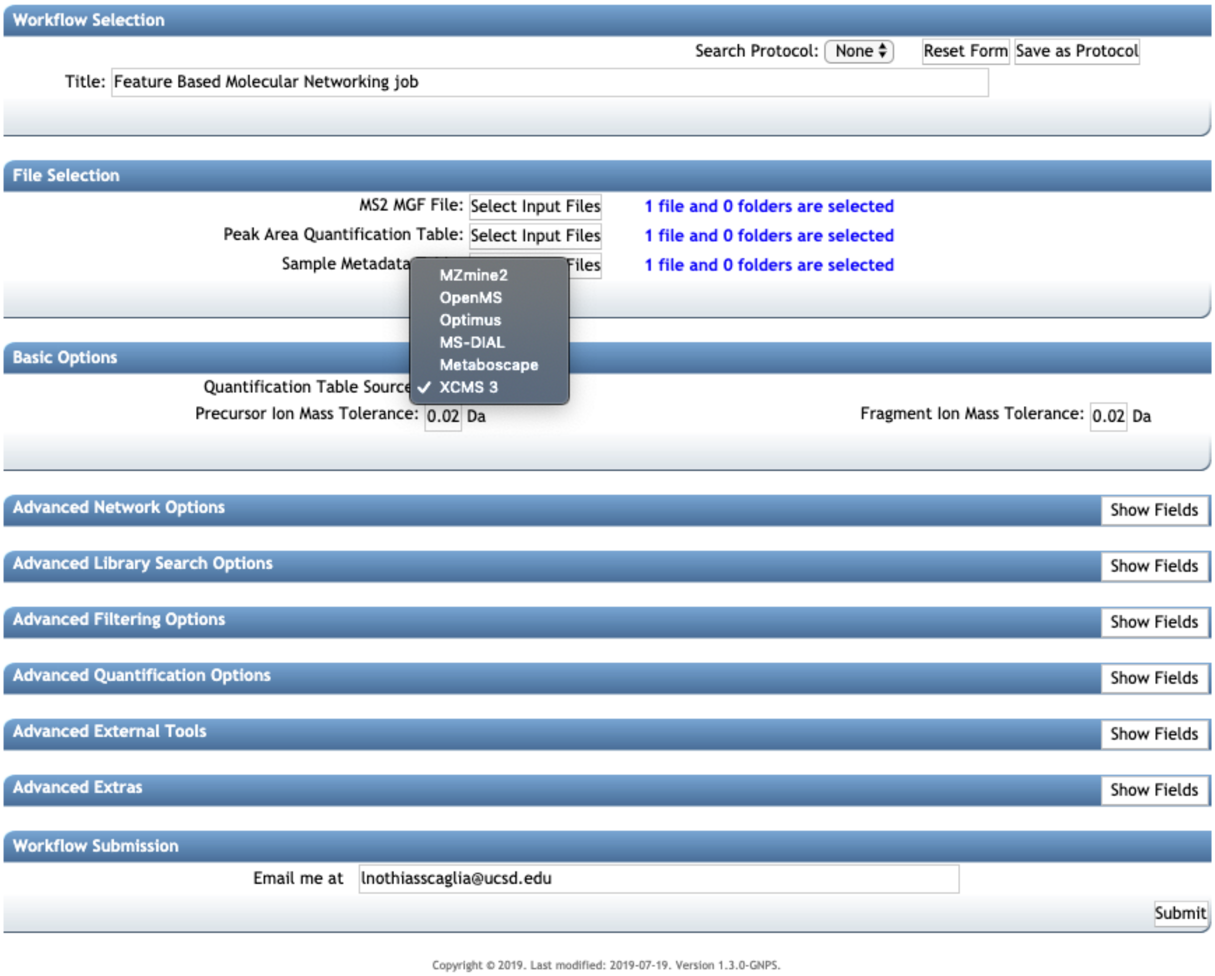


**Supplementary Figure 2.** Feature-based molecular networking workflow standard interface on GNPS (https://gnps.ucsd.edu).

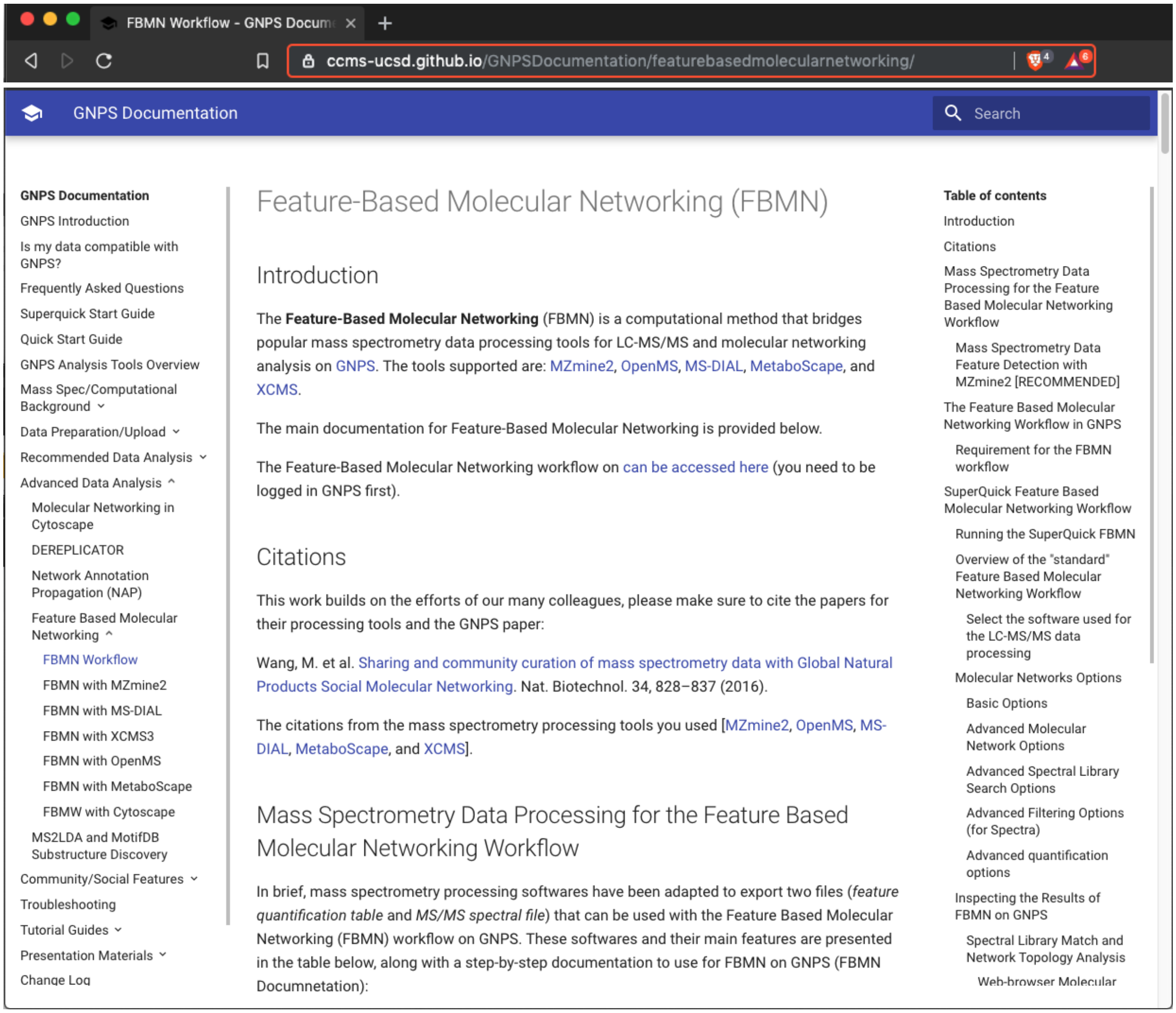


**Supplementary Figure 3.** The feature-based molecular networking documentation on GNPS (**https://ccms-ucsd.github.io/GNPSDocumentation/featurebasedmolecularnetworking/**).

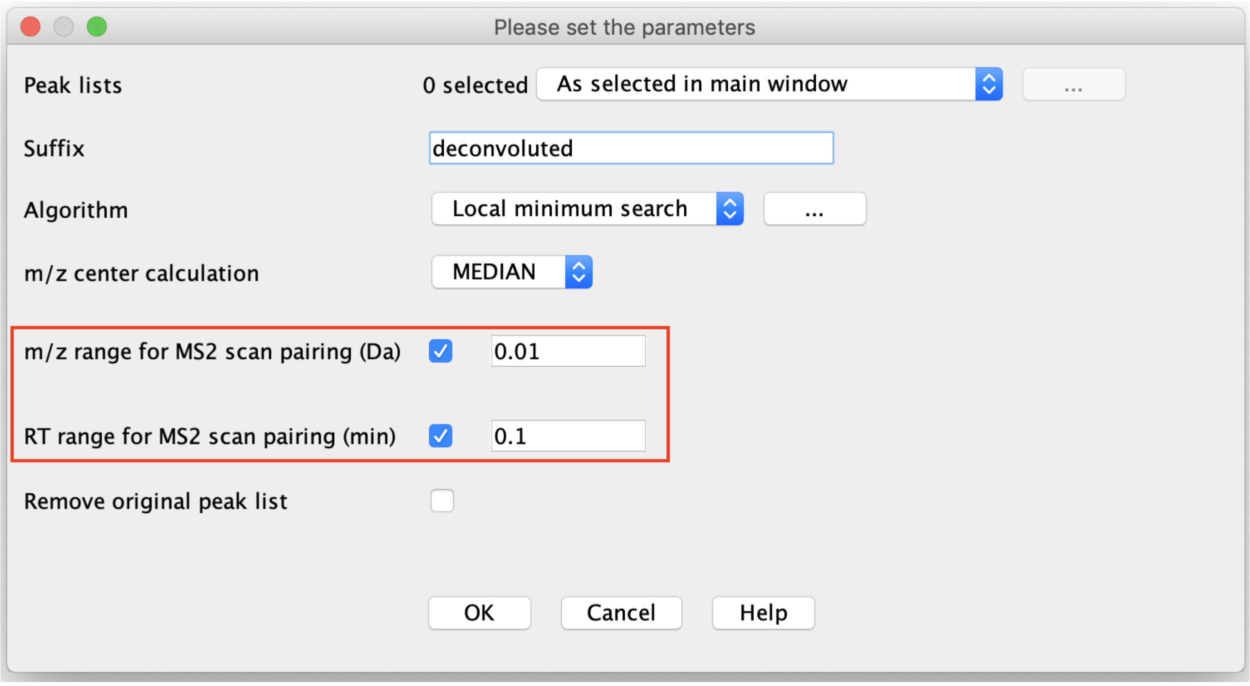


**Supplementary Figure 4.** The *Chromatogram deconvolution* module in MZmine and the options added for the pairing of $MS^1$ feature and $MS^2$ scans available since MZmine 2.27.

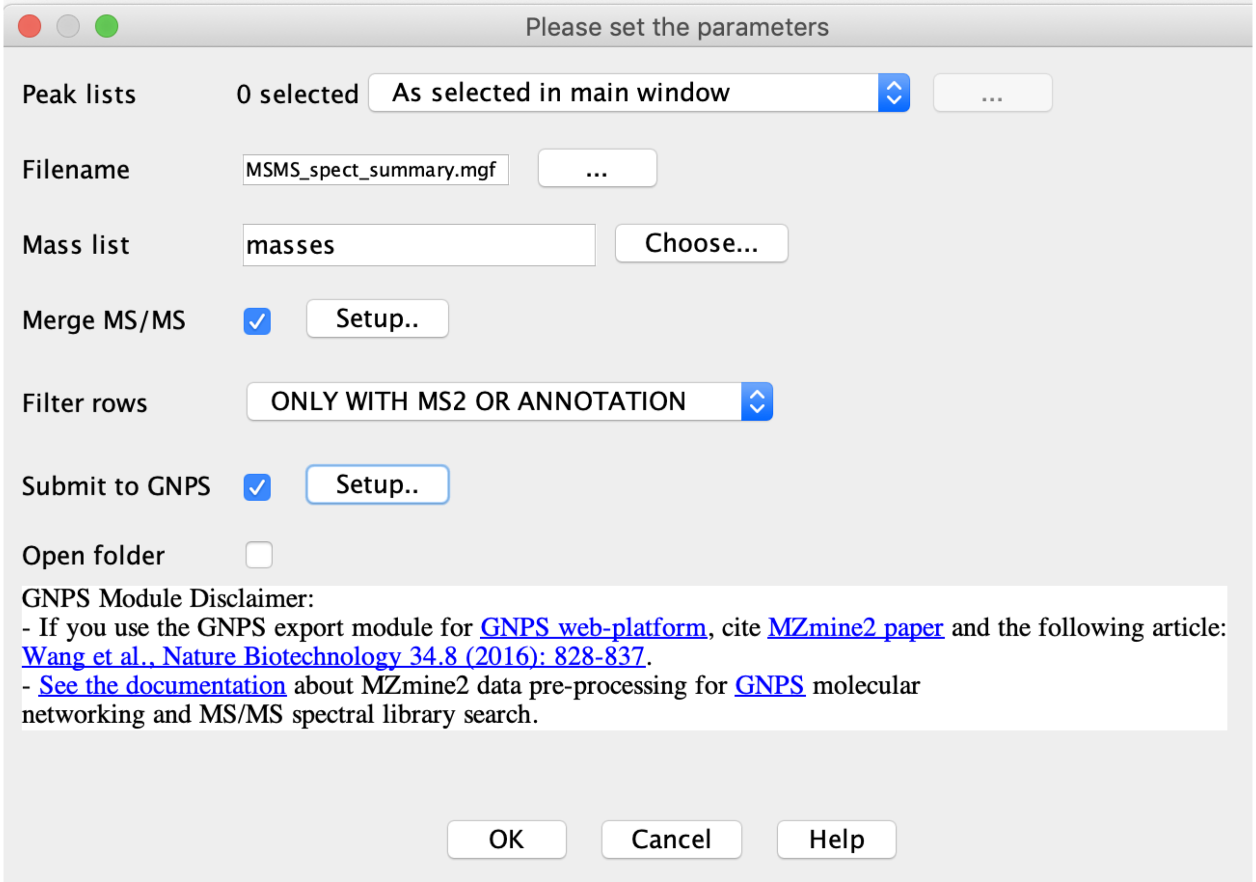


**Supplementary Figure 5.** The *Export for/Submit to GNPS* module interface in MZmine, available since version MZmine 2.37.

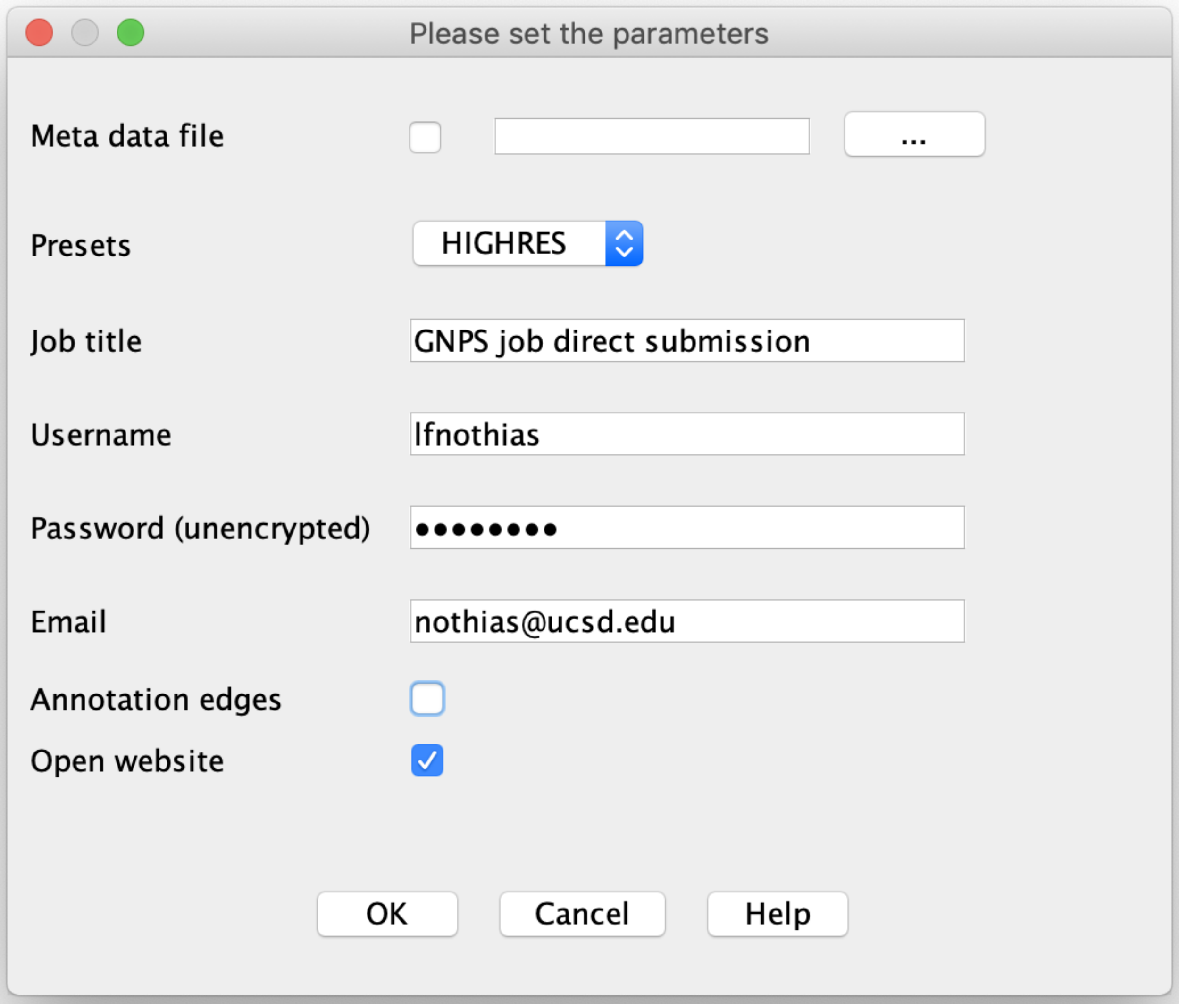


**Supplementary Figure 6.** Direct submission to GNPS with the *Export for/Submit to GNPS* module in MZmine.

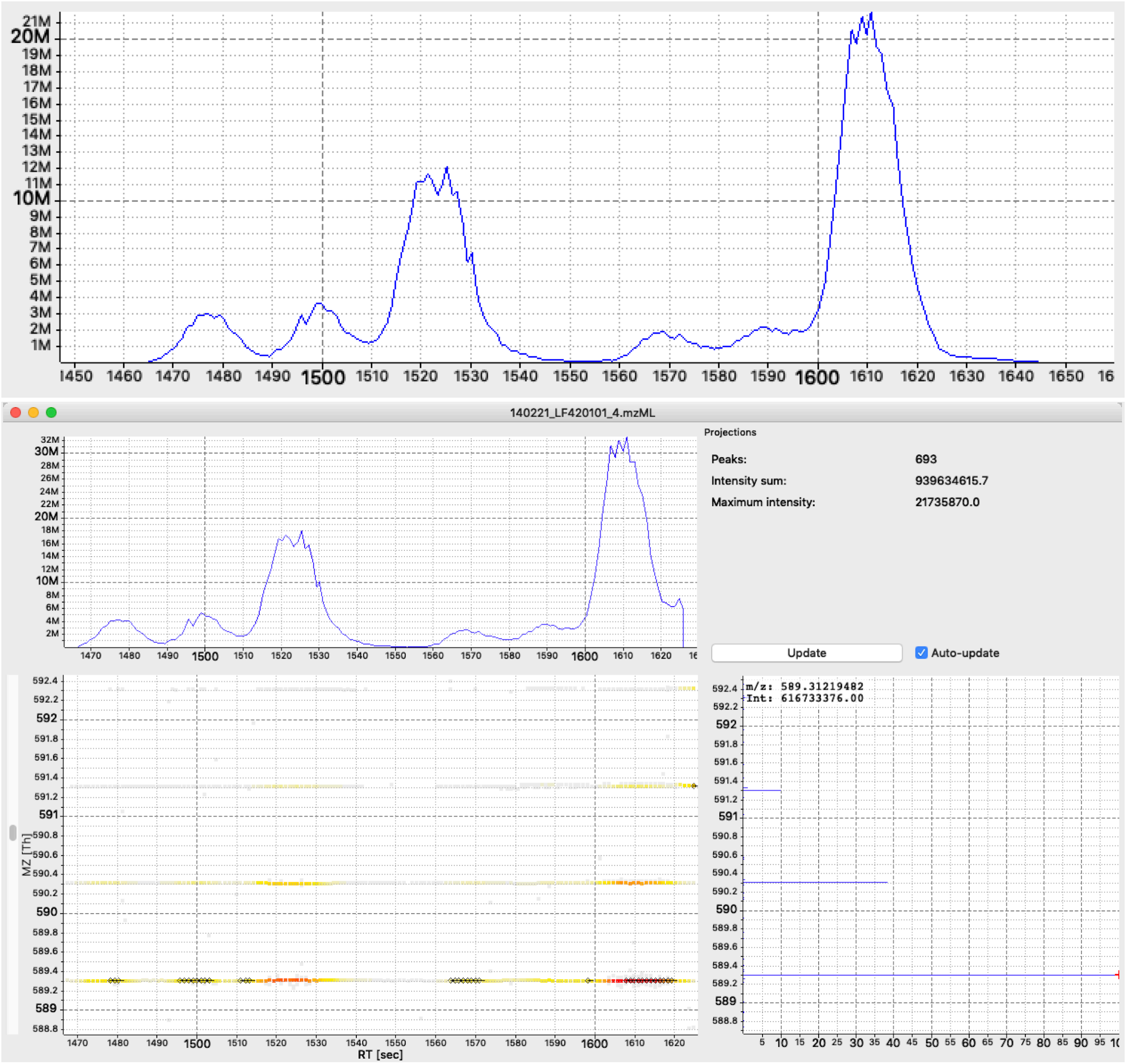


**Supplementary Figure 7.** Screenshots from OpenMS TOPPView. Top: Extraction Ion Chromatogram (EIC) of *m*/*z* 589.31 for the *Euphorbia dendroides* extract (LF420101_4.mzML) in the range 1450-1650 seconds. Bottom: Two dimensional LC-MS view of the compounds the isotopic pattern for the *m*/*z* 589.31 for the extract of *Euphorbia dendroides*. The top panel shows the EIC for the range *m*/*z* 589-592 and 1470-1640 seconds. The lower left panel shows the different intensities per *m*/*z* values, and the presence of fragmentation scans (black dots). The right panel shows the full spectrum for the range.

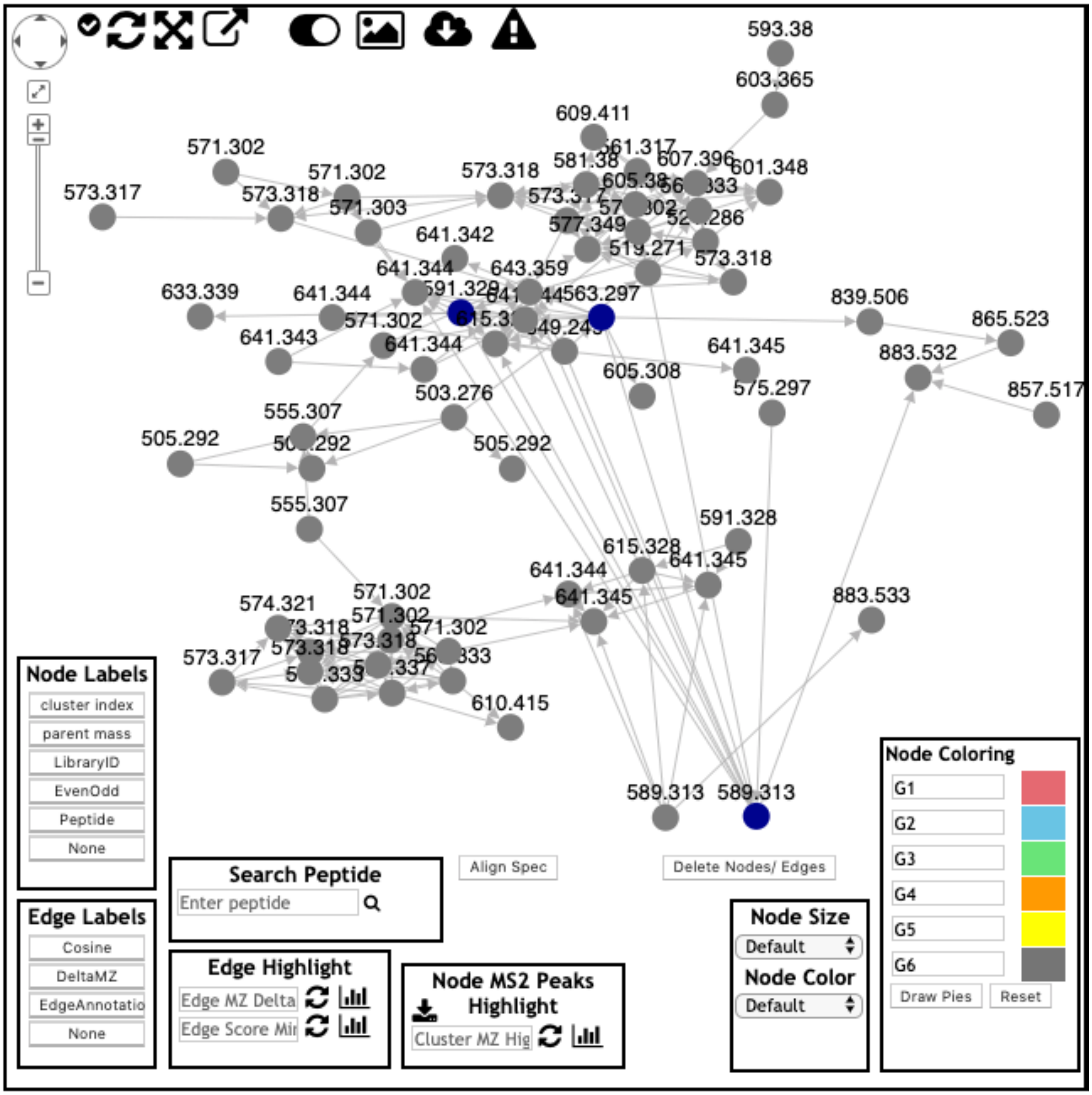


**Supplementary Figure 8.** Classical molecular networking analysis of *Euphorbia dendroides* dataset. View of the deoxyphorbol ester molecular network.

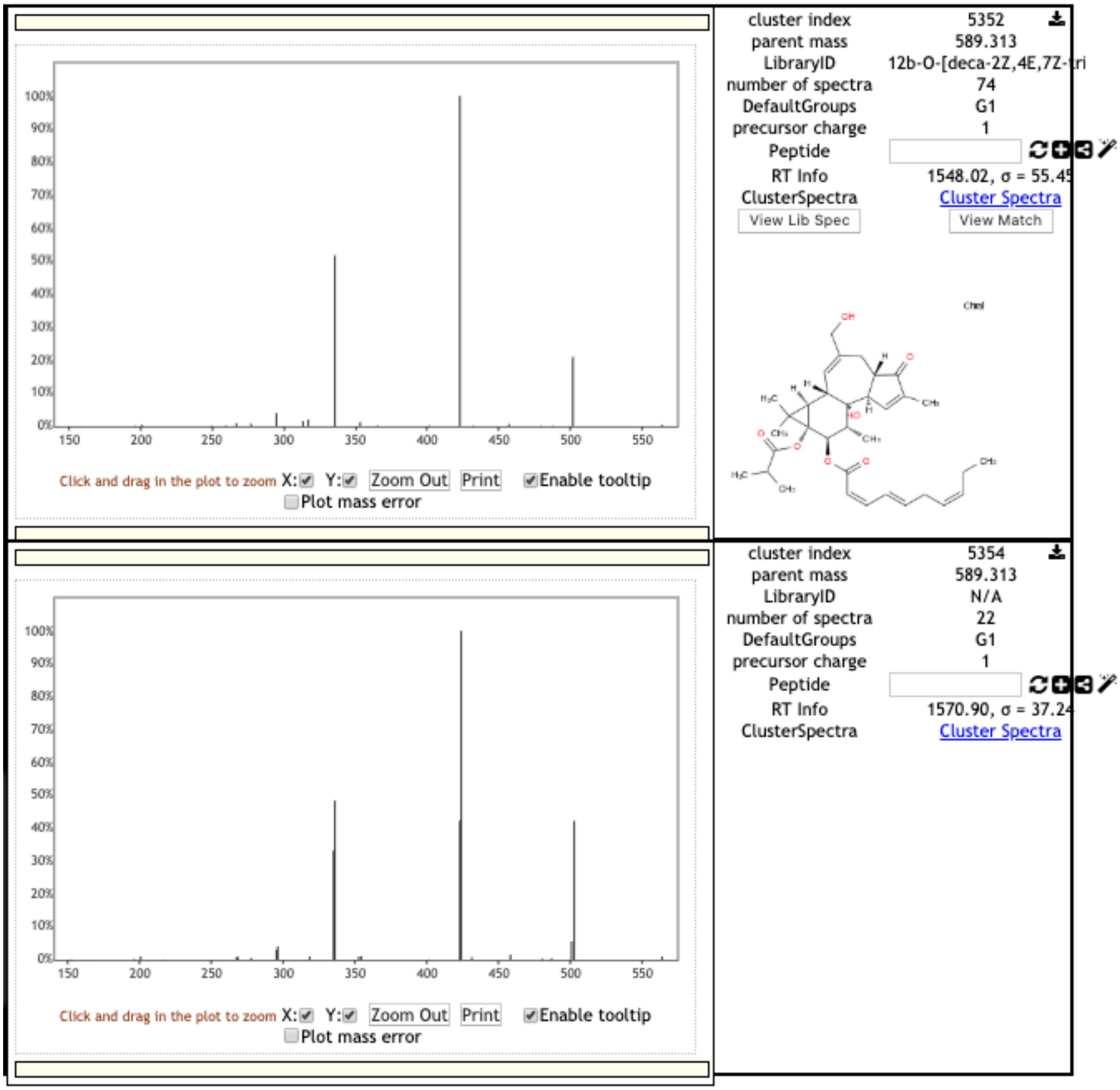


**Supplementary Figure 9.** Classical molecular networking analysis of *Euphorbia dendroides* dataset. View of the $MS^2$ spectra for the two at *m/z* 589.313 nodes.

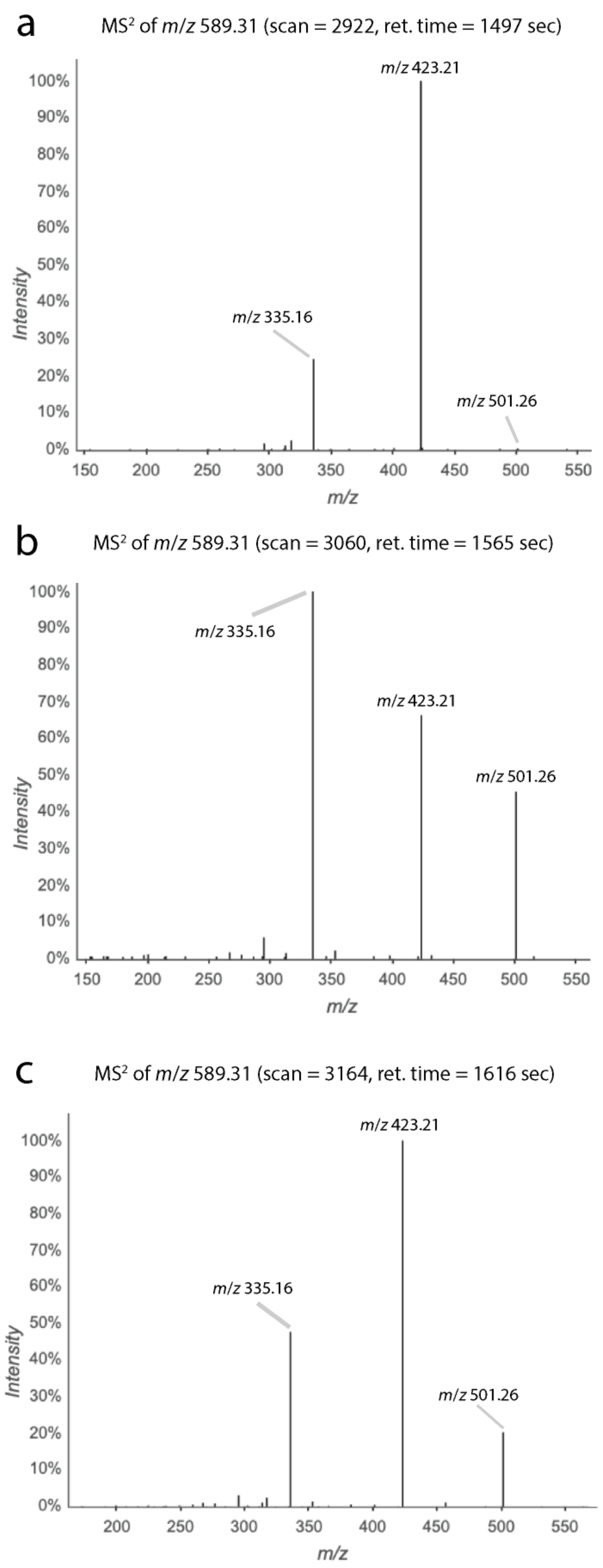
a
MS² of m/z 589.31 (scan = 2922, ret. time = 1497 sec)
m/z 423.21
m/z 335.16
m/z 501.26
Intensity
m/z
b
MS² of m/z 589.31 (scan = 3060, ret. time = 1565 sec)
m/z 335.16
m/z 423.21
m/z 501.26
Intensity
m/z
c
MS² of m/z 589.31 (scan = 3164, ret. time = 1616 sec)
m/z 423.21
m/z 335.16
m/z 501.26
Intensity
m/z

**Supplementary Figure 10**. Different $MS^2$ spectra for the ion *m/z* 589.31 observed in the *E. dendroides* extract. (a) The first spectral type was observed for the peaks at 24.6 - 25.2 min and characterized by a base peak at *m/z* 423.21 and a fragment ion *m/z* 335.16 with 20% relative intensity. (b) The second spectral type was observed for peaks at 26.1 min and a base peak at *m/z* 335.16, and fragment ions *m/z* 423.21 and 501.26 with 70-80% and 50-60% of relative intensities, respectively. (c) The third population was found for the chromatographic peaks at 26.1 - 27.0 min with a base peak at *m/z* 423.21 and fragment ions *m/z* 335.16 and 501.26 with 35-45% and 15-25% of relative intensities, respectively.

**Supplementary Note 2: Detailed discussion about the differences observed between classical molecular networking vs FBMN methods for *Euphorbia dendroides* data.**

A comparison of results between classical molecular networking vs FBMN (with MZmine) for the *Euphorbia dendroides* dataset is presented in Supplementary Table 1. FBMN is heavily dependent upon user-defined parameters selected during all steps of processing, including peak picking, chromatogram building and deconvolution, isotope grouping, feature alignment and gap filling. The discussion here described differences between the results obtained with classical and FBMN. For the *E. dendroides* dataset processed in MZmine with parameters described in the method, we observed that classical molecular networking produced more network nodes than FBMN. However, FBMN offered a higher spectral annotation rate (5.23%) than classical molecular networking (2.16% with a minimum cluster size = 2). The ratio between unique annotations/total annotations showed that FBMN is capable of separating different isomers, which get merged into one spectrum through the MS-Cluster algorithm in the classical molecular networking workflow (0.42 instead of 0.78 for classical molecular networking). However FBMN was found to provide less unique annotations than classical molecular networking (17 versus 22). This may indicate that relevant features were filtered out during MZmine processing for the FBMN using selected parameters.

MZmine offers various heuristic filters that can be used to reduce the number of features detected. We investigated the results of these filters on the network topology and annotation. When the filters “minimum 2 isotopes detected” and “minimum 2 occurrences” were used for FBMN analysis, the number of nodes became 315 nodes instead of 765 when no filter is used. Moreover, these two filters decreased the proportion of single nodes (31.4% instead of 44.8%), and increased the spectral annotation rate (10.47% instead of 5.23%) which suggests that they efficiently removed low quality spectra. Nevertheless, the use of these filters in MZmine decreased the number of nodes in the molecular network analysis, as seen in the deoxyphorbol molecular family (19 versus 39 nodes without filters). This can be explained by the nature of this dataset, which contains unique spectra from fractions of an *E. dendroides* extract, likely exhibiting high chemical diversity with many unique compounds. Thus, the use of the “minimum 2 occurrences” filter in MZmine, is filtering them out. In addition, the use of the filter “2 minimum isotopes detected” will filter out features that were detected at the limit of detection, and for which no $C_{13}$ isotopic peak can be observed/paired.

As different parameters selected during MZmine processing affect the final outcome, also the use of different processing software might influence results retrieved. The present FBMN results with MZmine enabled the discrimination of more isomers for the ion *m/z* 589.311, than when OpenMS[5] was used in the original paper[6], which illustrates how different processing software could lead to different results based on the parameters used and their specificities.

**Supplementary Table 1.** Results for classical molecular networking and feature-based molecular networking (with MZmine) for the *Euphorbia dendroides* dataset.

| | **Number of nodes** | **Single nodes** | **Unique library annotations** | **Total library annotations** | **Spectral annotation rate** | **Size of the deoxyphorbol ester network** |
|---|---|---|---|---|---|---|
| Classical molecular networking (minimum cluster size = 2, default) | 1,297 | 519 (40.0%) | 22 | 28 | 2.16 % | 22 (538 total spectra) |
| Feature based molecular networking without feature filters | 765 | 342 (44.7%) | 17 | 40 | 5.23 % | 39 |
| Feature based molecular networking with feature filter ("minimum 2 isotopes detected", "minimum 2 occurrences") | 315 | 98 (31.1%) | 15 | 33 | 10.47% | 19 |

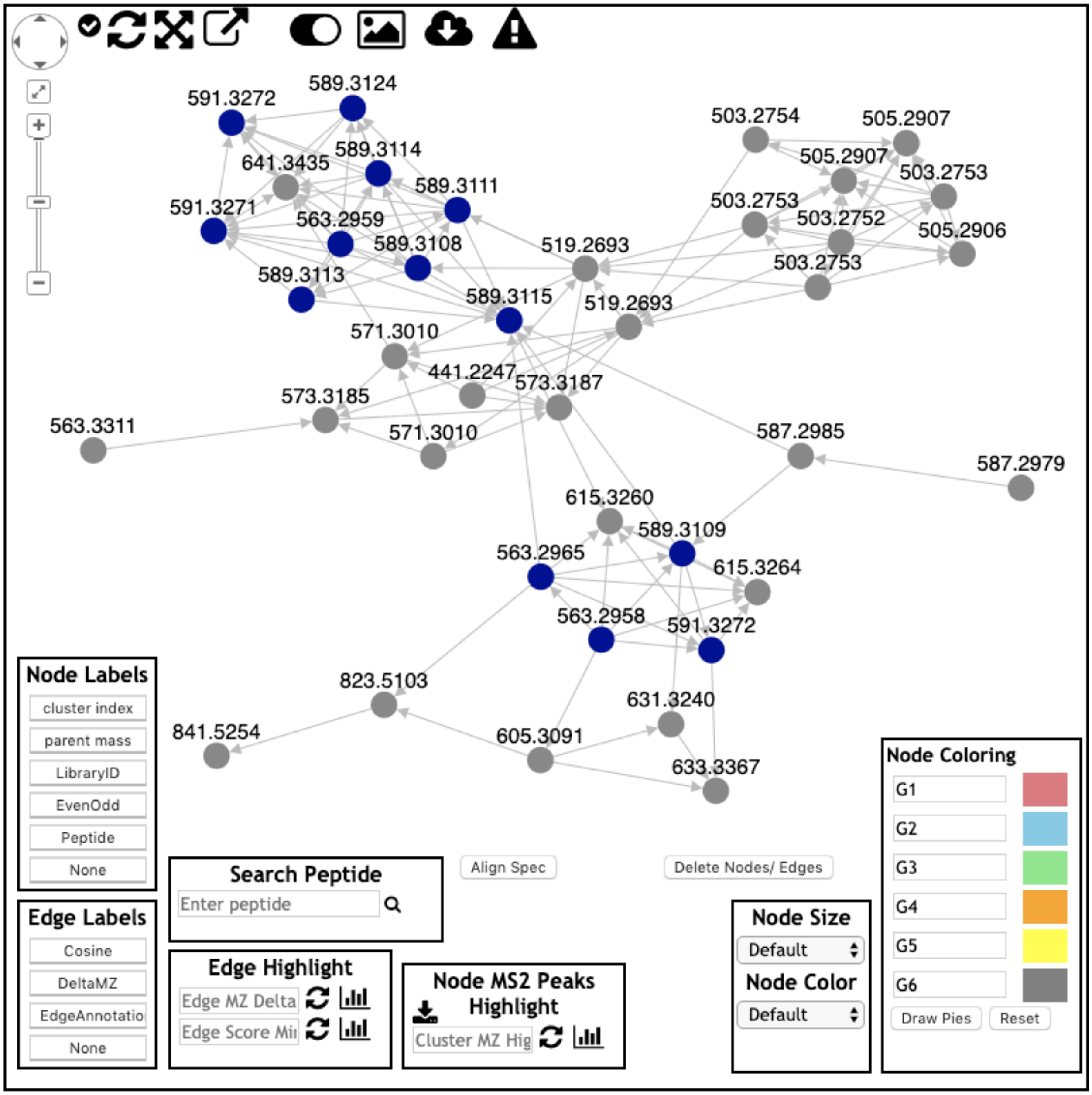


**Supplementary Figure 11**. Results of feature-based molecular networking with MZmine for the *E. dendroides* datasets. View of the entire deoxyphorbol cluster molecular network.

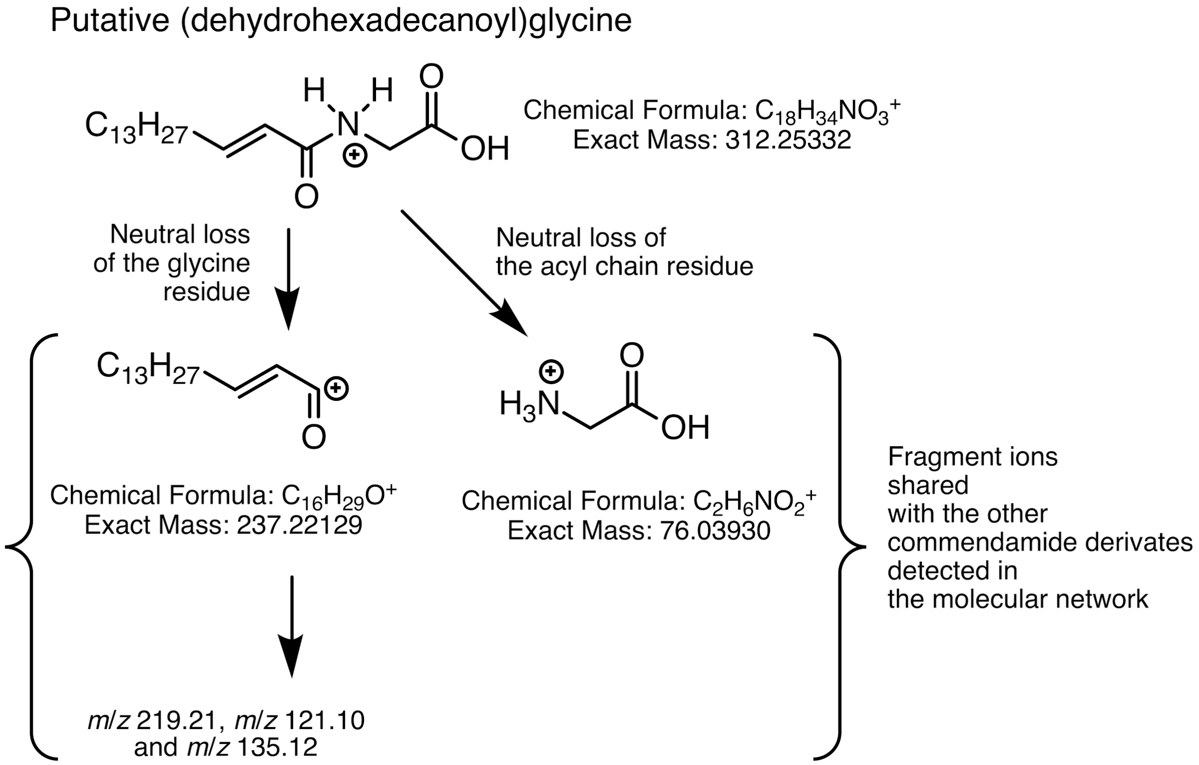


**Supplementary Figure 12**. Annotation of (dehydrohexadecanoyl)glycine, a putative commendamide derivative in the American Gut Project dataset (dehydrohexadecanoyl)glycine using FBMN on GNPS.

**Supplementary Note 3: Protocol for the MassIVE MSV00008263 dataset (EDTA case).**

**Sample preparation.** The 96-well plate PhreeTM Phospholipid Removal Kit was rinsed with 300 $\mu$L of MeOH (100%) and centrifuged at 500 g for 5 min, three times prior to sample addition. Blood plasma was stored at -80°C prior to extraction. The plasma microtubes were thawed at room temperature prior to extraction. Plasma samples were placed randomly into one of two PhreeTM Phospholipid Removal Kit 96-well plates. The thawed plasma samples were vortexed for 5 s and centrifuged for 1 min @ 5000 rpm prior to pipetting 50 µL of each sample into the 96-well PhreeTM Phospholipid Removal Kit. 200 µL of MeOH (100%) with 500 ng mL-1 lithocholic acid - d4 as an internal standard was added to each sample well using a multichannel pipette; the solution was aspirated and dispensed five times to mix plasma and organic solvent. 250 µL of a mixture of the bile acid standards at a concentration of 1000 ng mL-1, individually, in MeOH-Water (4:1) was added to 3 separate well (split 2 on one SPE plate and 1 on the other). A 96-well plate (Eppendorf® Microplate 96/U-PP) was placed under the PhreeTM Phospholipid Removal Kit to collect the sample, and centrifuged at 500 g for 5 min. The PhreeTM Phospholipid Removal Kit portion was discarded  and the sample-containing 96-well plate was evaporated until dry using a CentriVap Benchtop Vacuum Concentrator

(Labconco, Kansas City, MO, USA). The 96-well plate containing the dried extract were covered (Storage Mat IIITM 3080) and stored at -80°C prior to analysis. Immediately prior to analysis, the dried extract material was resuspended in 250 µL of MeOH-water (1:1) with 250 ng mL-1 cholic acid - d4, sonicated for 5 min, centrifuged for 5 min at 500 g, and covered with a plate sealing film (Zone-FreeTM Sealing Films).

**Data acquisition.** Plasma metabolite extracts were analyzed using an ultra-high performance liquid chromatography device (Vanquish, Thermo Fisher Scientific, MA, USA) coupled with an Orbitrap mass spectrometer (Q Exactive, Thermo Fisher Scientific, MA, USA). Chromatographic separation was carried out using a Kinetix C18 1.7 µm, 100 Å, 50 x 2.1 mm column with corresponding C18 guard cartridge maintained at 40°C during separation. 5.0 µL of extract was injected per sample. Mobile phase composition was as follows: A, water with 0.1% formic acid (v/v) and B, acetonitrile with 0.1% formic acid (v/v). Gradient elution was performed as follows: 0.0, 5.0% B; 1.0, 5.0% B; 1.1, 25.0% B; 5.0, 60.0% B; 5.75, 100.0% B; 6.5, 100%; 6.6, 5.0% B; 7.0, 5.0% B. Flow rate of 0.5 mL min-1 was held constant. Heated electrospray ionization (HESI) was performed in positive ion mode using the following source parameters: spray voltage, 3500 V; capillary temperature, 380 °C, sheath gas, 60.00 (a.u.); auxiliary gas, 20.00 (a.u.); sweep gas, 3.00 (a.u); probe temperature, 300 °C; and S-lens RF level, 60. The data-dependent acquisition parameters were set as follows: MS1 scans were collected at 30,000 resolution from m/z 150 to 1500 (~7 Hz) with a maximum injection time of 100 ms, 1 microscan, and an automatic gain control target of $1x10^6$. The top 3 most abundant precursor ions in the MS1 scan were selected for fragmentation with an m/z isolation width of 1.5 and subsequently fragmented with stepped normalized collision energy of 20, 30, and 40. The MS2 data was collected at 17,500 resolution with a maximum injection time of 100 ms, 1 microscan, and an automatic gain control target of $5x10^5$.

**Supplementary Note 4: FBMN makes it possible to achieve relative quantification**

**Sample Preparation**. The NIST SRM-1950 was prepared and extracted with 80% ethanol as proposed in the SRM 1950 paper[7].

**Mass Spectrometry Analysis**. The SRM1950 sample was analyzed using an ultra-high pressure liquid chromatography system (Vanquish, **Thermo** Fisher Scientific, MA, USA) coupled to an Orbitrap mass spectrometer (Q Exactive, **Thermo** Fisher Scientific, MA, USA) fitted with a heated electrospray ionization (HESI-II) probe. Chromatographic separation was accomplished using a Kinetex $C_{18}$ 1.3 µm, 100 Å, 2.1 mm x 50 mm column fitted with a C18 guard cartridge (Phenomenex) with a flow rate of 0.5 mL/min. 5 µL of extract was injected per sample/QC. The column compartment and autosampler were held at 40°C and 4°C respectively throughout all runs. Mobile phase composition was: A, LC-MS grade water with 0.1 % formic acid (v/v) and B, LC-MS grade acetonitrile with 0.1 % formic acid (v/v). The chromatographic elution gradient was: 0.0 - 1.0 min, 5% B; 1.0 - 9.0 min, 100% B; 9.0 - 11.0 min, 100% B; 11.0 - 11.5 min, 5% B; and 11.5 - 12.5 min, 5% B. Heated electrospray ionization parameters were: spray voltage, 3.5 kV; capillary temperature, 380.0 °C; sheath gas flow rate,

60.0 (arb. units); auxiliary gas flow rate, 20.0 (a. u.); auxiliary gas heater temperature, 300.0 °C; and S-lens RF, 60 (arb. units). MS data was acquired in positive mode using a data dependent method with a resolution of 35,000 in MS1 and a resolution of 17,000 in MS2. An MS1 scan from 100-1500 m/z was followed by an MS2 scan, using collision induced dissociation, of the five most abundant ions from the prior MS1 scan.

**Data interpretation.** Classical molecular networking does not use "area under the curve" the relative quantitative information obtained through feature detection of LC-MS traces. The FBMN method brings in ion abundance across all samples by using the value of chromatographic peak areas or peak heights as determined by the LC-MS feature finding software. Using a serial dilution of the NIST 1950 serum reference metabolome sample[7] analyzed on a Orbitrap instrument and process with OpenMS, we show the linearity of the relative quantification with FBMN and reveals the improvement compared to classical molecular networking (Figure 2g-h). As mentioned above, the most important limitation of classical molecular networking is the lack of dependable relative quantitative information. The interpretation of non-targeted LC-MS data in metabolomics relies on the statistical analysis of relative variation between ions intensity across the studied samples[8]. Classical molecular networking uses MS-Cluster results to define the ion distribution between the samples, using either the number of scans clustered in a consensus $MS^2$ spectrum ($MS^2$ scan table) or the sum of their precursor ion(s) intensities ($MS^2$ bucket table). A more accurate comparison of LC-MS data requires the value of chromatographic peak areas (EIC feature) or peak height for each ion features detected and aligned across the samples studied. The FBMN method makes possible to determine the relative intensity of each node in all samples. Using a serial dilution of the NIST 1950 serum reference metabolome sample analyzed on a Q Exactive mass spectrometer[9], we compared the capability of classical molecular networking and FBMN to evaluate the expected relative ion abundance. After basic optimisation of the software parameters, the use of MZmine and OpenMS resulted in feature intensities that correlated with high pearson correlation values for true positive compounds between the serial dilution with a median correlation of 0.92 (MZmine) and 0.73 (OpenMS), respectively. These differences observed between MZmine and OpenMS processing can be explained by the lack of a gap-filling step with OpenMS (See Supporting Informations) which results in less features detected in the low concentration range. However, most importantly, the results of classical molecular networking showed that neither the number of scans or the sum of precursor ion intensity were able to obtain satisfying correlation score ($r$ median 0.4 and 0.43) for the known compounds. The distribution of the $r$ value showed that 90% and 75% of the features had a value of 0.33 using the number of scans, and of the sum of the precursor ion intensity, respectively. The prevalence of that value in both metrics can be explained by the prevalence of features being selected for $MS^2$ scans in the most concentrated samples, but not in less concentrated samples. This shows that, while classical molecular networking can be used for qualitative analysis, it is not suited for accurate ion intensity statistical analysis and that binary metrics (presence/absence) might be better suited to use. Nevertheless, binary interpretation of classical molecular networking should also be considered with caution, as the absence of $MS^2$ spectra for a compound in one sample does not necessarily mean that the compound was not present, rather than simply below the signal threshold in order to selected for $MS^2$ in data-dependent acquisition (DDA) or absent due to other reasons.

**Supplementary Note 5: Combining FBMN with other mass spectrometry annotation tools.**

**SIRIUS**. SIRIUS is an advanced software for the computational annotation of small molecules from LC-MS/MS data. It is capable of annotating compounds at the molecular formula, structural and substructural level[10], and uses both isotopic pattern from $MS^1$ scans and the compound $MS^2$ spectra[11–13]. The *spectral* $MS^2$ *summary file* (MGF file format) generated for the FBMN is compatible with SIRIUS, and can provide putative or partial annotation of the molecular network, which is essential since spectral library matching usually results in 1-5% annotation rate. In addition, a *Sirius Export* module/function was created in MZmine and MetaboScape that exports representative $MS^1$/$MS^2$ spectra for each feature. The $MS^1$ spectra contains information about the detected isotopic pattern and can be used for automatic adduct/rare elements detection in SIRIUS[11], which restricts the molecular formula search space to speed up computation and improve molecular formula identification rates[13].

**DEREPLICATOR**. DEREPLICATOR[14], along with DEREPLICATOR VarQuest[15], are a collection of computational mass spectrometry tools specialized in the annotation peptidic small molecules often produced by micro-organisms endowed with various biological activities. DEREPLICATOR tools can be run directly through the FBMN workflow results on GNPS. Alternatively and for advanced parametring, the DEREPLICATOR workflow on GNPS accepts the *$MS^2$ spectral summary* file (.MGF format) as input and can directly map into the feature-based molecular networks (https://ccms-ucsd.github.io/GNPSDocumentation/dereplicator/).

**Network Annotation Propagation.** Network Annotation Propagation (NAP)[16] uses MetFrag/MetFusion[17,18] for the prediction of putative structures, and utilizes the network topology to rerank structure predictions by propagating the expected structural similarity. NAP is available on GNPS as a dedicated workflow, and offers direct support to FBMN (https://ccms-ucsd.github.io/GNPSDocumentation/nap/).

**Unsupervised substructure annotation with MS2LDA.** MS2LDA uses the Latent Dirichlet Allocation algorithm to mine for motifs (Mass2Motifs) of co-occurring fragments and neutral losses in the $MS^2$ spectra.[19,20] MS2LDA accepts the *$MS^2$ spectral summary file* used in feature-based molecular networking, therefore the mapping between MS2LDA results and the network can be performed directly on the GNPS web-platform (http://gnps.ucsd.edu), or using MolNetEnhancer[21], or a python script.[22] MS2LDA results produced on the GNPS web-platform can be exported for import into the MS2LDA visualisation tool (ms2lda.org).

**MolNetEnhancer.** MolNetEnhancer combines the outputs from molecular networking, MS2LDA, and *in silico* structure annotation tools (including Network Annotation Propagation and DEREPLICATOR) together with automated chemical classification through ClassyFire[23] into a single molecular network[21]. MolNetEnhancer accepts input files from classical as well as feature-based molecular networking and provides a comprehensive overview at the chemical class as well as substructure level. The MolNetEnhancer algorithm is publicly available on Github (https://github.com/madeleineernst/pyMolNetEnhancer, https://github.com/madeleineernst/RMolNetEnhancer) or through the GNPS web-platform (https://ccms-ucsd.github.io/GNPSDocumentation/molnetenhancer/).

**Supplementary Note 6: Large dataset processing for FBMN.**

The processing of large metabolomics dataset (more than a few thousand samples) is limited by the scalability of existing LC-MS feature detection tools, especially for those based on graphical user interfaces (such as MZmine and MS-DIAL). MZmine was successfully used for the processing of large datasets acquired on QTOF mass spectrometers, but required both a powerful workstation computer (with 64GB of RAM memory), and the use of sub-optimal parameters, such as higher noise thresholds, to reduce the computational load. Here, we show that the use of XCMS enables the processing of large metabolomics studies for FBMN analysis. The MassIVE dataset MSV000080030 consists of approximately 2,000 samples from the forensic study with samples from hands and objects of 80 participants analyzed on a QTOF by LC-MS/MS[24]. The files were processed with XCMS running on cluster computers. For XCMS, the processing was performed on 8 processors with 32 GB of RAM memory allocated for each process. The XCMS script is available at https://github.com/DorresteinLaboratory/XCMS3_FeatureBasedMN and the GNPS job (https://gnps.ucsd.edu/ProteoSAFe/status.jsp?task=cf026a37c70946a1a937e030dea65514). The number of features detected and spectral library matches show that, for large datasets FBMN will result in less annotations (0.7% instead of 1.0% for classical molecular networking, in Supplementary Table 2). Indeed, these feature detection tools were not designed to process multi-thousands files datasets. As a result, the processing of 2,000 files requires sub-optimal parameters (noise level, feature intensity threshold) to limit the computational load, and of heuristic(s) (such as a minimum number of occurrences in the dataset, or the presence of isotopologues) to limit the number of features outputted for downstream analysis. This illustrates the continued need for improvement of LC-MS feature detection on large metabolomics datasets, that will enable sensitive detection within a reasonable runtime.

**Supplementary Table 2.** Comparison of the results for the MassIVE MSV000080030 dataset (QTOF) including 2000 samples using classical molecular networking and feature-based molecular networking with XCMS.

| | Number of features | Single nodes | Unique library annotations | Spectral annotation rate | Runtime for molecular networkingon GNPS |
|---|---|---|---|---|---|
| Classical molecular networking | 20,578 (nodes) | 18,946 (92,1%) | 208 | 1.0 % | 11h44min |
| Feature-based molecular networking with XCMS | 13,862 (LC-MS features with MS2) | 12,045 (86.9%) | 97 | 0.7 % | 6h20min |

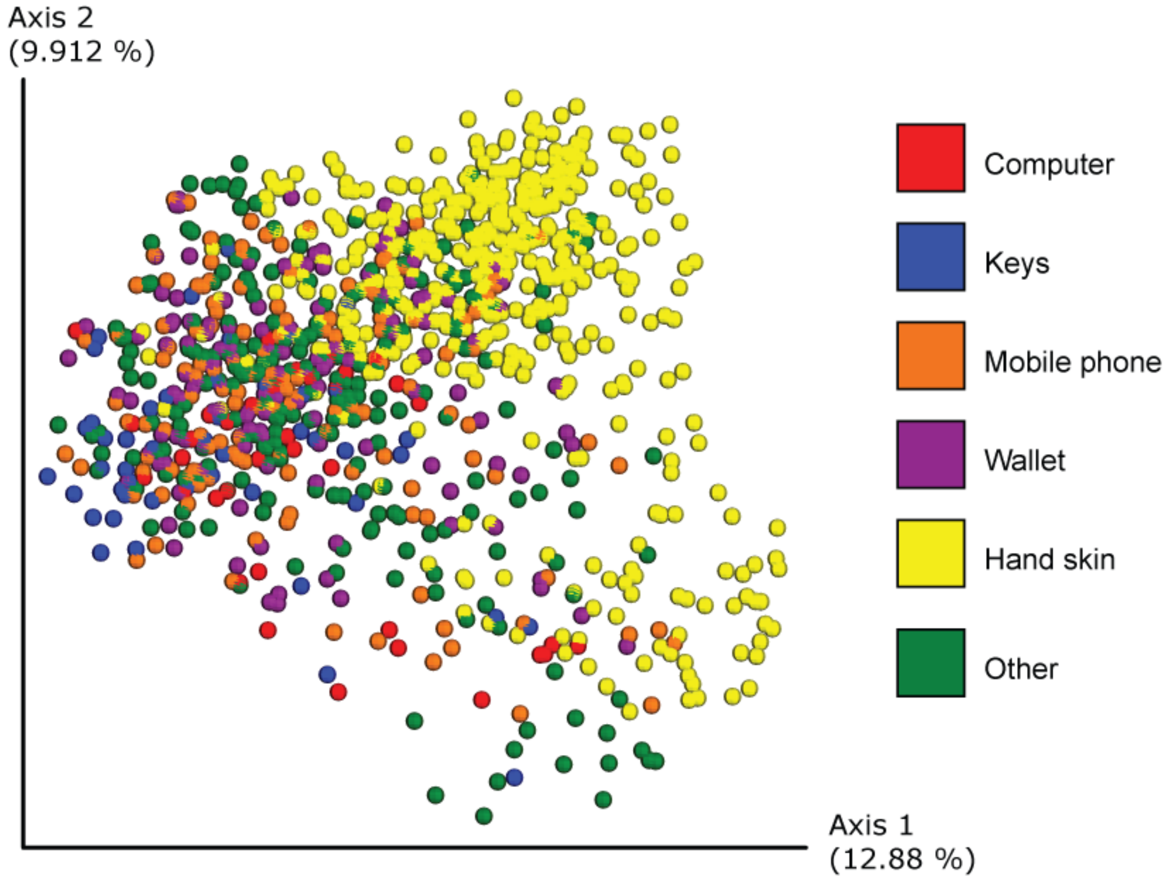


**Supplementary Figure 130**. Principal Coordinate Analysis of the forensic study (MSV000080030, approximatively 2,000 samples) processed with XCMS and analyzed with the FBMN workflow on GNPS. See this link for an interactive visualization of the plot.
